# Identification of neural progenitor cells and their progeny reveals long distance migration in the developing octopus brain

**DOI:** 10.1101/2021.03.29.437526

**Authors:** Astrid Deryckere, Ruth Styfhals, Ali Murat Elagoz, Gregory E. Maes, Eve Seuntjens

## Abstract

Cephalopods have evolved nervous systems that parallel the complexity of mammalian brains in terms of neuronal numbers and richness in behavioral output. How the cephalopod brain develops has only been described at the morphological level, and it remains unclear where the progenitor cells are located and what molecular factors drive neurogenesis. Using histological techniques, we located dividing cells, neural progenitors and postmitotic neurons in *Octopus vulgaris* embryos. Our results indicate that progenitors are located outside the central brain cords in the lateral lips adjacent to the eyes, suggesting that newly formed neurons migrate into the cords. Lineage tracing experiments then showed that progenitors, depending on their location in the lateral lips, generate neurons for the different lobes. The finding that octopus newborn neurons migrate over long distances is reminiscent of vertebrate neurogenesis and suggests it might be a fundamental strategy for large brain development.

## Introduction

Cephalopod mollusks represent an invertebrate lineage that exhibits morphological as well as behavioral complexity reminiscent of vertebrates. Studying species from this group thus brings an opportunity to understand the genetic drivers of the development of organ systems that evolved convergently with vertebrates. The adult cephalopod mollusk *Octopus vulgaris* has a highly centralized brain containing about 200 million nerve cells in the supra- and subesophageal mass and two optic lobes (Young, 1963, 1971), yet the cellular and molecular mechanisms driving brain development remain poorly understood. At hatching, the *O. vulgaris* brain counts about 200,000 cells and occupies roughly one fourth of the total body, indicating extensive embryonic neurogenesis (Budelmann, 1995; Giuditta et al., 1971; Packard & Albergoni, 1970). In general, neural progenitor cells are generated from ectodermal cells and divide symmetrically and asymmetrically to generate all neurons of the nervous system (Florio & Huttner, 2014; Kriegstein & Alvarez-Buylla, 2009; Wodarz & Huttner, 2003). In clades harboring species with diffuse nerve nets such as cnidaria and hemichordates, the proliferating neural progenitor cells are distributed throughout the ectoderm generating local neurons, while in (sub)phyla with a centralized nervous system including vertebrates, arthropods and some annelids, the neural progenitor cells are grouped in the neurectoderm. The proliferating neural progenitor cells either remain on the apical surface of the neurectoderm in vertebrates and annelids, or internalize as is the case for insects (Cunningham & Casey, 2014; Götz & Huttner, 2005; Hartenstein & Stollewerk, 2015; Lowe et al., 2003; Meyer & Seaver, 2009; Rentzsch et al., 2017; Simionato et al., 2008; Taverna et al., 2014; Urbach & Technau, 2004). In addition, long-distance migration of neurons has been described for developing vertebrate brains, in which neurons born in different zones follow longer trajectories to their final location, where they intermingle to form complex circuits (García-Moreno & Molnár, 2020). Although neuronal migration has been described in developing invertebrate nervous systems as well, this process is generally limited to restricted cell populations, e.g. the Q, CAN and HSN neuroblasts in *Caenorhabditis elegans* (Blelloch et al., 1999; Forrester et al., 1998; Montell, 1999), or to short-range migratory events, e.g. in the *Drosophila* visual system (Apitz & Salecker, 2015; Bhat, 2007; Morante et al., 2011).

Molecular studies in vertebrates, *Drosophila*, *Caenorhabditis elegans*, but also cnidarians, have revealed a set of regulatory transcription factors involved in neurogenesis (Arendt et al., 2008; Bertrand et al., 2002; Galliot et al., 2009; Hirth, 2010; Layden et al., 2012). First, group B SRY-related HMG box genes (*soxB* genes) regulate the generation of the neurectoderm and also maintain neurectodermal cells in an undifferentiated and proliferative state (Guth & Wegner, 2008; Sarkar & Hochedlinger, 2013). After neurectoderm formation, members of the superfamily of basic helix-loop-helix (bHLH) transcription factors control the specification of neural progenitor cells and also activate neuronal differentiation pathways (Bertrand et al., 2002; Vervoort & Ledent, 2001). The bHLH protein superfamily is divided in groups A-F, based on structural and biochemical properties. bHLH genes such as *atonal*, *neurogenic differentiation* (*neuroD*) and *neurogenin* (*ngn*), and *achaete-scute* (*asc*) complex members are classified in group A. The role of bHLH genes is best described in nervous system development where members of this group A (and some of group B) regulate the patterning, differentiation and specification of neurons, from sponges to primates (Powell & Jarman, 2012; Simionato et al., 2007; Vervoort & Ledent, 2001). Although the level of differentiation of progenitor cells in which these factors are expressed and the sequential expression of bHLH genes differs slightly across species, they all steer progenitor cells towards a neural fate. After this neural progenitor commitment, regulatory genes such as *musashi*, *prospero*, and *embryonic lethal abnormal vision* (*elav*) (vertebrate *hu*), will activate programs to initiate differentiation of progenitor cells into neurons (Choksi et al., 2006; Okano et al., 2002; Pascale et al., 2008). In the end, postmitotic neurons will mature, form synapses, and produce neurotransmitters to form a fully functional central nervous system (CNS).

These neurogenic processes are poorly studied in lophotrochozoans, and above all, in lophotrochozoans with a complex nervous system such as *O. vulgaris*. How and where neurons are generated, whether they migrate and what factors drive their differentiation and integration remain unknown. The high fecundity of *O. vulgaris*, spawning small and transparent eggs that can be kept in a remote tank system, as well as the available genomic data recently made it a tractable species for embryonic studies (Deryckere et al., 2020; Zarrella et al., 2019). Here, we have visualized the anatomy of brain development in 3D using light sheet microscopy. Expression analysis of several transcription factors, proliferation and neural markers revealed the spatial and temporal pattern of *O. vulgaris* brain development from stage IX onwards. Lastly, lineage tracing studies using CFDA-SE flash labeling proved that the neurogenic area is spatially patterned. Our data suggest that *O. vulgaris* deploys a neurogenic transcription factor sequence during neurogenesis, as well as extensive neural migration, that are reminiscent of vertebrate mechanisms of large brain generation.

## Results

### The rapidly growing octopus brain suggests massive embryonic neurogenesis

*O. vulgaris* displays a direct embryonic development, giving rise to actively feeding paralarvae. Their embryonic development takes approximately 40 days at 19 °C and has been classified in 20 major stages I-XX, with some stages subdivided in an early and late part (Deryckere et al., 2020; Naef, 1928). Organogenesis starts at Stage VII.2 and can be split in early, mid- and late phases (Figure 1A). In the early organogenesis phase, the brain anlagen (cordal) are elongated and interconnected (in 3 dimensions) (Figure 1A,B) as also described by Shigeno et al. for *Octopus bimaculoides* (Shigeno et al., 2015). On the most anterior side, lateral and posterior to the mouth and foregut, the cerebral cord (CC) will give rise to the lobes of the supraesophageal mass (SEM), while on the posterior side of the embryo, the palliovisceral (PVC) and pedal (PC) cords will form the subesophageal mass (SUB). The two bilateral optic cords (OC) will differentiate and grow to form the optic lobes (OL) (Figure 1A,C). At hatching, the central brain that now surrounds the esophagus is considerably big and contains densely packed nuclei (Figure 1D). The brain is not yet fully mature, but the major lobes described for the sub- and supraesophageal masses can be distinguished (Figure 1E,F) (Young, 1971). In contrast to the growing brain cords, the tissue adjacent to the eye primordia first grows in size from Stage VII.2 to Stage XV.2, and then shrinks to disappear at hatching, suggesting this tissue might contribute to the inner head structures (Figure 1A). This tissue was first described as “Kopflappen” by Marquis and later as “anterior chamber organ” by Shigeno et al., Yamamoto et al. and Koenig et al. (Koenig et al., 2016; Marquis, 1989; Shigeno et al., 2001; Yamamoto et al., 2003). However, the term “anterior chamber organ” was introduced by Young, referring to neurovenous tissue adjacent to the adult eye, that presumably regulates the fluid content between the lens and cornea (Young, 1970). The term was most likely wrongly adopted embryonically as the tissue at that stage is not restricted to the anterior side of each eye, but is rather placed lateral to the eye fields. The structure is also connected to the central brain, through a stream-like transition zone. Because of its position and shape, we renamed the tissue to “lateral lips” (LL, indicated in mint green in Figure 1G) and introduced the term “posterior transition zone” (PTZ) for the region that interconnects the lateral lips with the central brain on the posterior side of the embryo (Figure 1H).

**Figure 1.**
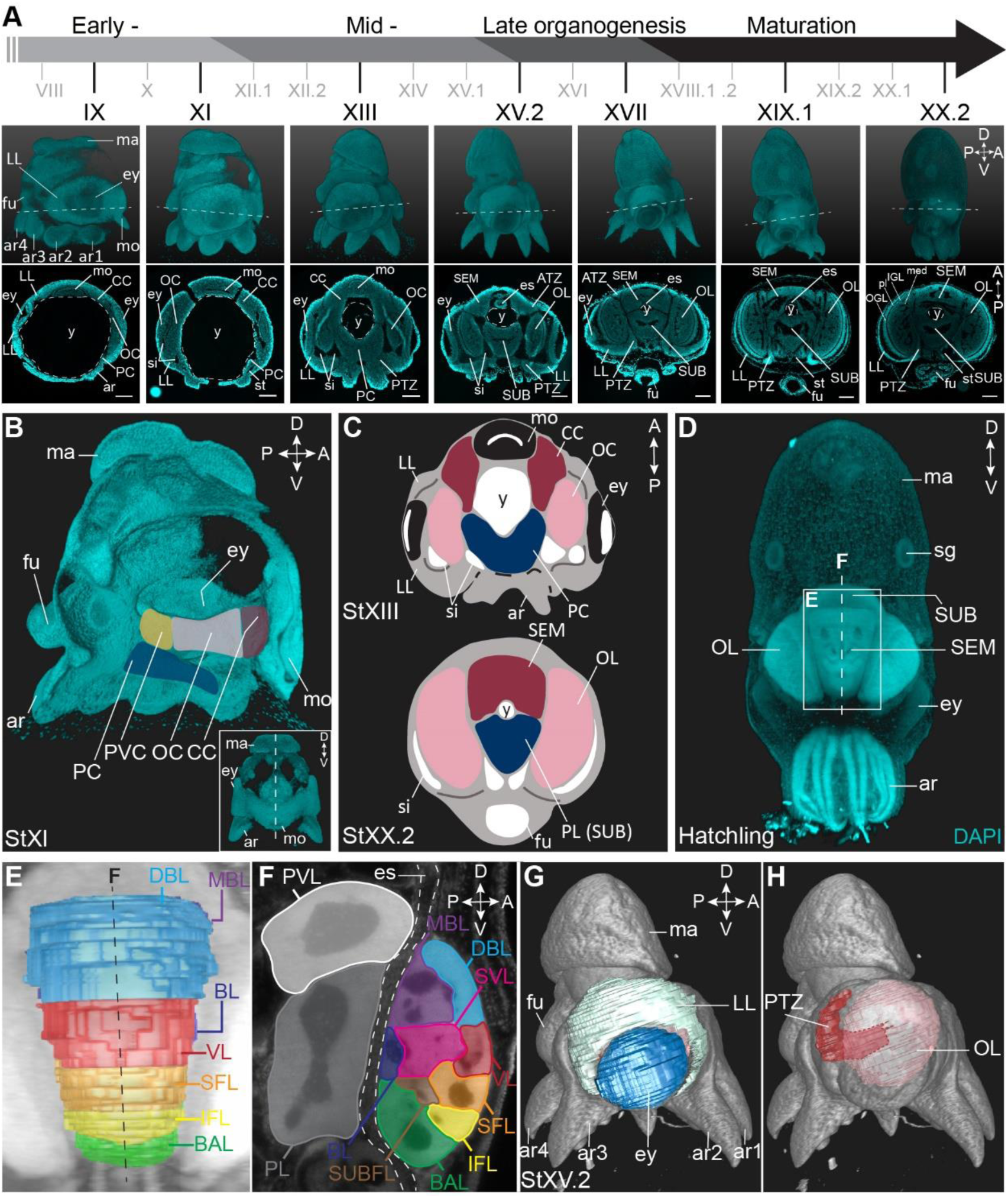
Overview of the developing *O. vulgaris* embryo and its nervous system. A. Overview of *O. vulgaris* embryonic development from Stage IX to Stage XX.2, covering early-, mid- and late-organogenesis events and maturation. Surface renderings after DAPI staining are shown in the upper panels (lateral view, dorsal side up) and representative transversal sections in the lower panels (anterior side up). The OCs that give rise to the OLs develop medially from the eye primordia. The CC that generates the SEM develops next to the external mouth and the PC and PVC that give rise to the SUB develop on the posterior side. The dashed lines on the surface renderings indicate the sectioning plane of the transversal sections. Scale bars represent 100 μm. B. Surface rendering after DAPI staining at Stage XI, showing that the central brain cords are connected and encircle the yolk. Prospective cords are pseudo-colored: CC in red, OC in pink, PVC in yellow and PC in blue. C. Schematic of the octopus head region mid-organogenesis and at hatching (OC/OL in pink, CC/SEM in red, PC/PL of the SUB in blue). D. Maximum projection after DAPI staining of a hatchling showing the densely nucleated central brain from the anterior side. E-F. Reconstruction of the different lobes in the SEM and SUB of a hatchling in 3D (E) and on an optical section (F). G-H. 3D reconstruction of the eye (blue), LL (mint green), OL (pink) and the PTZ (red) in a Stage XV.2 embryo (DAPI in grey). *Abbreviations: A, anterior; ar, arm; ATZ, anterior transition zone; BAL, buccal lobe; BL, basal lobe; CC, cerebral cord; D, dorsal; DBL, dorsal basal lobe; es, esophagus; ey, eye; fu, funnel; IFL, inferior frontal lobe; IGL, inner granular layer; LL, lateral lips; ma, mantle; MBL, medial basal lobe; med, medulla; mo, mouth; OC, optic cord; OGL, outer granular layer; OL, optic lobe; P, posterior; PC, pedal cord; pl, plexiform layer; PL, pedal lobe; PTZ, posterior transition zone; PVC, palliovisceral cord; PVL, palliovisceral lobe; SEM, supraesophageal mass; SFL, superior frontal lobe; sg, stellate ganglion; si, sinus ophthalmicus; st, statocyst; SUB, subesophageal mass; SUBFL, subfrontal lobe; SVL, subvertical lobe; V, ventral; VL, vertical lobe; y, yolk*.

### The developing octopus brain shows early neuronal differentiation

In order to map the (early) patterns of neuronal development in *O. vulgaris*, we studied the expression of the pan-neuronal *elav* gene. ELAV/Hu RNA-binding proteins are a family of splicing factors, predominantly present in differentiating neurons, from the moment they exit the cell cycle (Colombrita et al., 2013). Through blast searches against the full-length transcriptome of *O. vulgaris* embryos and paralarval brains generated in this study (see Materials and Methods), we identified three candidate *elav* transcripts. We performed a phylogenetic analysis on these ELAV proteins together with 27 ELAV sequences from 20 other species, 5 non-neural ELAV sequences from 4 other species and 5 PolyA-binding sequences in order to root the tree (Figure S2.1). One of the *O. vulgaris* ELAV candidate proteins nested with vertebrate HuC/D and invertebrate neural ELAV homologs. We refer to this sequence as *Ov-*ELAV. The two other candidate proteins could be identified as non-neural (Figure S2.1). We also performed a conserved domain (CD)-Search and detected an ELAV/HuD family splicing factor domain which is characteristic of ELAV proteins.

We mapped the expression of *Ov-elav* using *in situ* hybridization at embryonic stages IX, XI, XIII, XV.2, XVII, XIX.1 and XX.2 (Figure 2, S2.2). These stages serve as a starting point for the characterization of nervous system development during organogenesis (Stage IX-XVII) and maturation phases (Stage XIX.1 and XX.2) of embryonic development. *Ov-elav* transcripts were detected throughout development with little expression at Stage IX in the cerebral, palliovisceral, pedal and optic cords. Staining intensity increased by Stage XI, showing low-level staining in the lateral lips surrounding the eye placode and more intense staining in all cords and surrounding the mouth. A similar pattern was observed at subsequent stages with highest staining intensity in the central brain cords/masses and low intensity staining in the lateral lips. At Stage XIII, a region in between the lateral lips and the central brain on the posterior side of the embryo showed intermediate *Ov-elav* expression. This region corresponds to the posterior transition zone (PTZ), introduced earlier. From Stage XV.2 onwards, cells on the anterior side of the embryo and laterally from the supraesophageal mass, showed intermediate *Ov-elav* expression levels as well. Accordingly, this zone will be named the anterior transition zone (ATZ). In the optic lobes at stages XIX.1 and XX.2, *Ov-elav* expression was highest in the medulla and inner granular layer, but lower in the outer granular layer. The ATZ and PTZ shrink towards the end of embryonic development and seem to become integrated in the developing brain. In summary, our data indicate that neuronal differentiation is present at early organogenesis phases already, and the transition zones as well as the growing cords and masses contain large numbers of postmitotic neurons.

**Figure 2.**
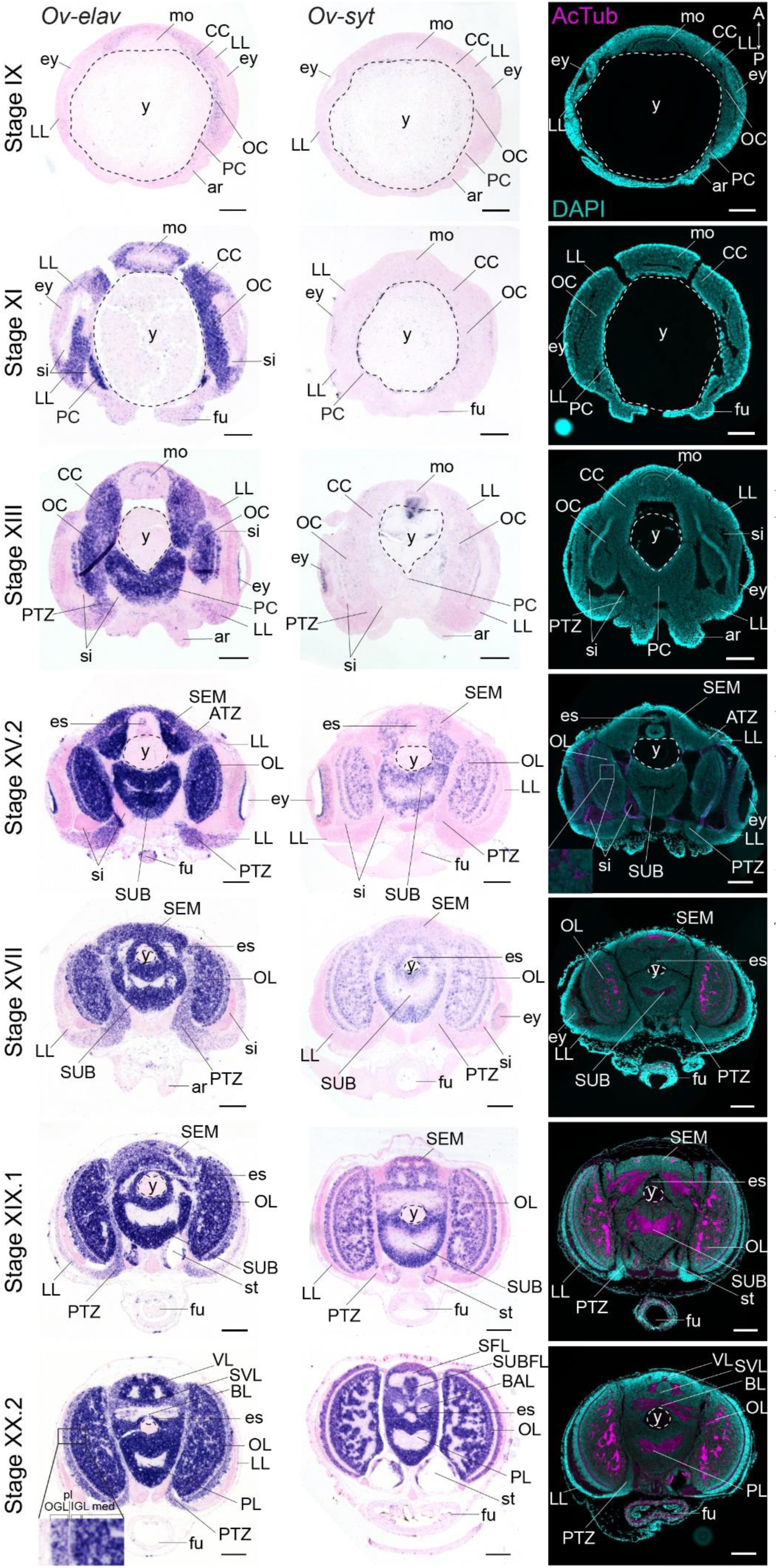
Formation of the *O. vulgaris* brain. *In situ* hybridization of *Ov-elav* (left column) and *Ov-syt* (middle column) and immunoreactivity against acetylated alpha tubulin (right column) on transversal sections of embryos at Stage IX, XI, XIII, XV.2, XVII, XIX.1 and XX.2 with anterior up and posterior down. *Ov-elav* expression levels are clearly elevated from Stage XI onwards and are generally highest in the developing brain cords and central brain masses, intermediate in the transition zones and low in the lateral lips. *Ov-syt* can be detected first in the retina at Stage XI and in the outer layers of the optic cord at Stage XIII. From Stage XV.2 onwards, transcripts are present in all brain masses. Acetylated alpha-tubulin is present in the optic lobes from Stage XV.2 onwards (low level, magnified box) and in the supra- and subesophageal masses from Stage XVII onwards. Scale bars represent 100 μm. *Abbreviations as in* Figure 1.

To address the maturation of these neurons, we investigated the expression pattern of *synaptotagmin*. Synaptotagmin is a presynaptic calcium sensor necessary for neurotransmitter release, present in mature synapses and thus in differentiated, functional neurons (Poskanzer et al., 2003). In addition, we also visualized neuronal somata and neuropil using immunoreactivity against acetylated alpha-tubulin, as previously described for multiple cephalopod species (Figure 2, S2.3) (Jung et al., 2018; Kingston et al., 2015; Scaros et al., 2018; Shigeno et al., 2015; Shigeno & Yamamoto, 2002; Wollesen et al., 2009, 2012). In *O. vulgaris*, we observed expression of *Ov-syt* in the retina from Stage XI onwards. In the central brain, it was first expressed in the outer layers of the optic cord at Stage XIII, and in the optic lobe medulla, supra- and subesophageal masses from Stage XV.2 onwards, pointing to a sequential maturation. First low-level immunoreactivity against acetylated alpha tubulin was found from Stage XV.2 onwards within the optic lobes. We therefore used the adult terminology (optic lobe, supra- and subesophageal mass) for CNS annotation from this stage onwards, since the embryonic cords demonstrate clear signs of differentiation.

### The lateral lips harbor the neurogenic zone whereas the developing cords are postmitotic

The developing octopus brain thus consists mainly of *Ov-elav* expressing cells, and it remains unclear where these cells are initially generated. A previous report in the squid *Doryteuthis pealeii* suggested that cells surrounding the eye placode contribute to the brain (Koenig et al., 2016). This area might therefore harbor the neurogenic zone during embryonic development. In order to locate proliferating cells that might contribute to CNS development, we used immunoreactivity against phospho-histone H3 (PH3) and mapped the expression pattern of *Ov-pcna*.

PCNA is a nuclear protein required for DNA replication and repair, and functions as a cofactor of DNA polymerase-delta in eukaryotes and archaea (Barry & Bell, 2006; Baserga, 1991; Prelich et al., 1987). Expression of *pcna* is tightly regulated and peaks at late G_1_ and S phases of the cell cycle (Santos et al., 2015). The expression of the *O. vulgaris* homolog was mapped using *in situ* hybridization at embryonic Stages VII.2, IX, XI, XIII, XV.2, XVII, XIX.1 and XX.2. *Ov-pcna* transcripts were detected throughout embryonic development (Figure 3). While at Stages VII.2 and IX, transcripts were found in most embryonic tissues (mouth apparatus, retina, lateral lips and cords, in the latter at lower intensity), transcripts gradually disappeared in the optic, cerebral, palliovisceral and pedal cords from Stage XI onwards. At Stage XI, most *Ov-pcna* expressing cells that were found adjacent to the eye, located to an area that is *Ov-elav* negative and that might be part of the epithelium lining the sinus ophthalmicus. *Ov-pcna* transcripts were dispersed throughout the lateral lips and were absent from the transition zones and central brain at Stages XV.2-XX.2. Since transcripts of *Ov-pcna* might still be present at low level in cells that finished DNA replication, we also mapped the phosphorylation of histone H3 during embryonic development (Figure 3). Histone H3 is phosphorylated on serine 10 and serine 28 during early mitosis and dephosphorylated at the end of mitosis in eukaryotes (Hans & Dimitrov, 2001; Prigent & Dimitrov, 2003; Wei et al., 1998). Immunohistochemistry on embryonic tissue from Stage VII.2 to XX.2 showed that mitotic activity in the embryonic head region was high at the beginning of organogenesis, with the highest level of dividing cells present in the mouth apparatus, at the apical side in the developing retina and spread throughout the lateral lips. Consistent with *Ov-pcna* expression, the number of PH3 positive cells in the cerebral, optic, palliovisceral and pedal cords was limited, with possibly very few, single, dim and seemingly randomly organized positive cells present until Stage XIII. At late organogenesis and maturation stages, PH3 positive cells located to the lateral lips and were absent from the central brain. Given the *elav*-positive areas are almost devoid of dividing cells, and the transition zone represents an area connecting the mitotically active lateral lips to the growing CNS, our data strongly suggest that the lateral lips represent the octopus neurogenic zone.

**Figure 3.**
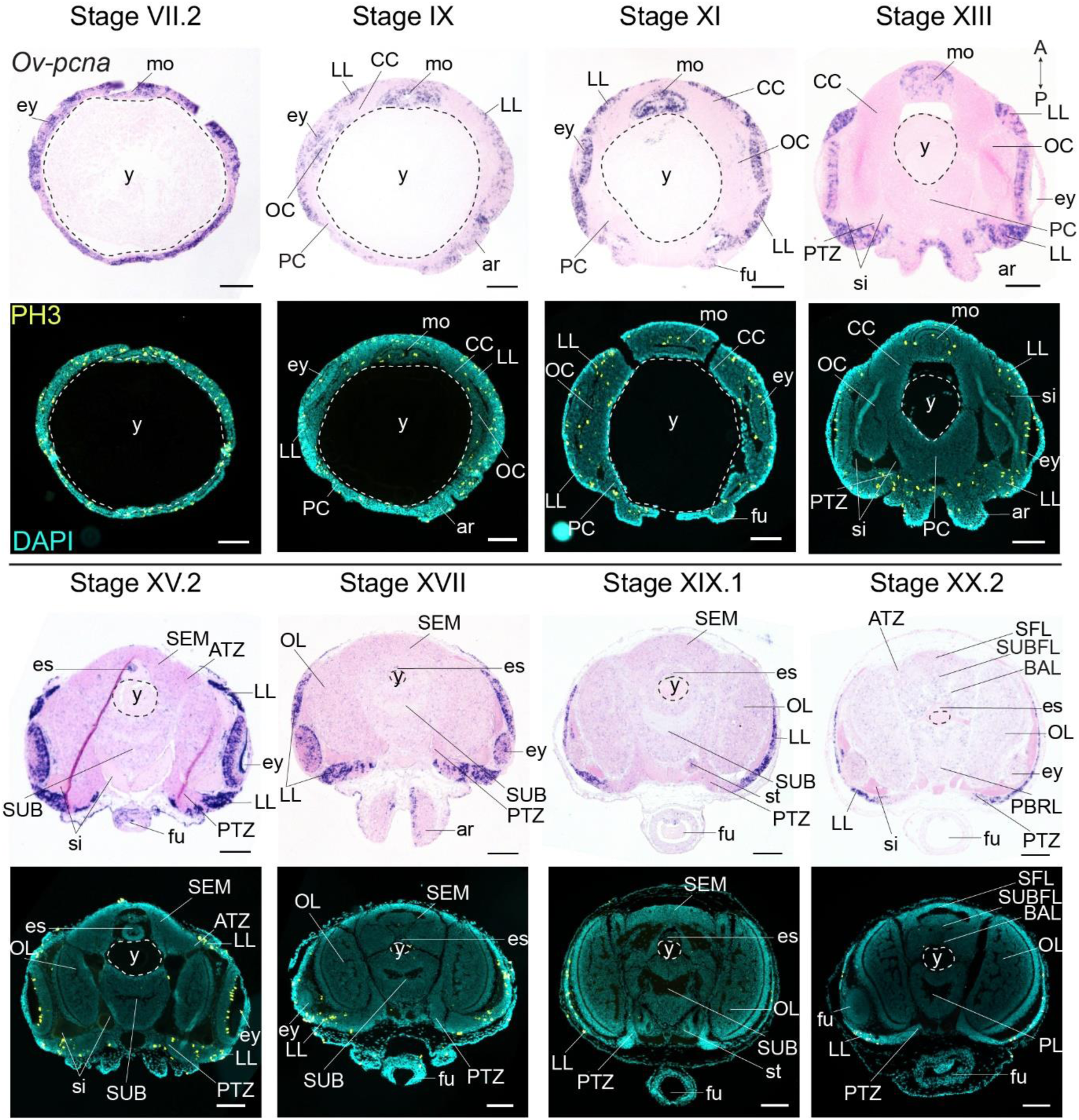
Cell division in the developing *O. vulgaris* embryo. Expression of *Ov-pcna* (upper panels) and immunoreactivity against PH3 (lower panels) from Stage VII.2 to Stage XX.2. *Ov-pcna* expression is broad at Stages VII.2 and IX and gets restricted to the lateral lips, mouth region and retina at subsequent stages. Similarly, PH3 positive cells are abundant in the lateral lips, in the mouth region and on apical side in the retina. Very few cells in the developing central brain are PH3 positive or express *Ov-pcna*. Scale bars represent 100 μm. *Abbreviations as in* Figure 1.

### Conserved sequence of neurogenic transcription factor gene expression during *O. vulgaris* brain development

In order to find molecular support for the neurogenic character of the lateral lip cells, we mapped the expression of conserved genes involved in neural stem cell specification and differentiation, namely *soxB1*, *ascl1*, *ngn* and *neuroD*.

First, to spatially map the extent of the neurectoderm, we studied the expression of *soxB1*. *Sox* genes have been divided into 7 groups (A-G) based on the sequence of their high mobility group (HMG)-box that binds to DNA in a sequence-specific manner (Sasai, 2001; Wegner, 1999). Members of the *SoxB* group play important roles in (early) neurogenesis in various bilaterians and can be further divided in two sub-groups SoxB1 and SoxB2 based on sequence differences outside the HMG-box and on their activity (Bowles et al., 2000; She & Yang, 2015). Using BLASTn and tBLASTn searches, we identified *O. vulgaris* members for groups B-E, but not for group A which is mammalian specific or group G which is also restricted to particular vertebrate lineages (Bowles et al., 2000; Heenan et al., 2016). We could also not identify SOXF family members in the available cephalopod transcriptomes of *O. vulgaris*, O. *bimaculoides*, *Sepia officinalis* and *Euprymna scolopes* (Figure S4.1). Based on phylogenetic analysis including 31 SOXB, 3 SOXC, 4 SOXD, 4 SOXE and 3 SOXF amino acid sequences from other species together with 5 TCF/LEF sequences to root the tree, we identified single *O. vulgaris* homologs for SOXC, D and E and two SOXB homologs (Figure S4.1). One *O. vulgaris* SOXB protein nested within the SOXB2 clade and the other one within the SOXB1 clade, the latter named *Ov-*SOXB1. CD-Search of this *Ov-*SOXB1 protein revealed the highly conserved SOX-TCF HMG-box and SOXp superfamily domains.

*Ov-soxB1* transcript levels were high in the head region at all embryonic stages examined (Figure S4.2). At Stage IX, *Ov-soxB1* was expressed in the lateral lips, eyes and surface ectoderm surrounding the mouth apparatus (esophagus, radula, salivary gland). In the developing central brain, *Ov-soxB1* was expressed in the cerebral cord, but not in the palliovisceral, pedal and optic cords. At Stage XI, *Ov-soxB1* was still highly expressed in the lateral lips and cerebral cord, and transcripts were now also present in the optic cord in a patched pattern. At Stage XIII, *Ov-soxB1* transcripts were most highly expressed in the cerebral cord and showed a similar patched pattern in the optic cords. Transcripts were also present in the pedal cord and limited in the palliovisceral cord. At Stage XV.2, *Ov-soxB1* expression was high in the lateral lips and supraesophageal mass. In the subesophageal mass, *Ov-soxB1* positive cells were more spread and in the optic lobes, *Ov-soxB1* transcripts showed the highest expression in the inner granular layer, while expression was more distributed in the outer granular layer and medulla. At Stage XVII, *Ov-soxB1* transcripts were still numerous in the central brain, which has started to form neuropil. At Stages XIX.1 and XX.2, *Ov-soxB1* was expressed in the lateral lips, the supra- and subesophageal mass and optic lobes where it was most highly expressed in the inner granular layer. Taken together, *Ov-soxB1* was not only expressed in putative stem cells, but also in postmitotic (*Ov-elav*^+^) areas, as has been described for other invertebrate species.

To identify a transcription factor that would mark neural progenitors only, we mapped the expression of two proneuronal bHLH group A transcription factors *ascl1* and *neurogenin*, and one neuronal differentiation bHLH transcription factor *neuroD*. By sequence similarity searches, we identified *O. vulgaris* homologs for *achaete-scute*, *neurogenin* and *neuroD*. Phylogenetic analysis of the atonal-related bHLH transcription factors including 15 NEUROD and 8 NEUROGENIN/TAP protein sequences from other species effectively assigned *Ov-neuroD* and *Ov-ngn* to their respective subfamilies (Figure S4.3). bHLH proteins possess a bHLH domain which is involved in DNA binding and dimerization (Jones, 2004; Murre et al., 1989). CD-Search identified such conserved domains in both sequences. In addition, phylogenetic analysis for the achaete-scute-related bHLH gene *Ov-ascl1* together with 27 other ASCa protein sequences and 10 ASCb protein sequences placed *Ov-*ASCL1 within the proneuronal ASCa subgroup of the bHLH superfamily (Figure S4.4). CD-Search revealed a highly conserved DNA-binding bHLH domain.

Achaete-scute homologs are involved in vertebrate neural identity determination and are key regulators in *Drosophila* neuroblast generation (Cabrera et al., 1987; Skeath & Carroll, 1992; Vervoort & Ledent, 2001). In *O. vulgaris*, *Ov-ascl1* transcripts were detected throughout embryonic development at all stages tested (Figure 4, S4.5). At Stage IX, *Ov-ascl1* was expressed in the lateral lips surrounding the eye primordia. Expression could also be observed in retinal cells of the developing eyes and in the tissue delineating the mouth apparatus, and was absent from the cerebral, pedal, palliovisceral and optic cords. At subsequent stages of organogenesis (XI, XIII, XV.2 and XVII), *Ov-ascl1* was expressed in the lateral lips and was absent from the developing cords/brain lobes. During maturation and right before hatching, the number of cells expressing *Ov-ascl1* significantly decreased with the thinning lateral lips and transcripts were still absent from the optic lobes, supra- and subesophageal masses. Similar to *achaete-scute*, homologs of *neurogenin* are highly conserved proneural genes expressed in early nervous system development, before neuronal differentiation in vertebrates, but also annelids (Simionato et al., 2007, 2008; Sur et al., 2017; Vervoort & Ledent, 2001). In *O. vulgaris* embryos, we found *Ov-ngn* transcripts in the lateral lips throughout organogenesis and maturation phases, while transcripts were absent from the developing brain cords (Stage IX-XX.2, Figure 4, S4.6). In addition, very few *Ov-ngn* transcripts were present in both transition zones at Stages XIII to XX.2. Equivalent to *Ov-ascl1*, the number of *Ov-ngn* expressing cells decreased considerably in the maturation phase with the thinning of the lateral lips and at Stage XX.2, only few cells were expressing *Ov-ngn*. In summary, expression of both *Ov-ascl1* and *Ov-ngn* consistently marks the proliferative lateral lips, with *Ov-ascl1* expressing cells being more numerous compared to *Ov-ngn* expressing cells.

**Figure 4.**
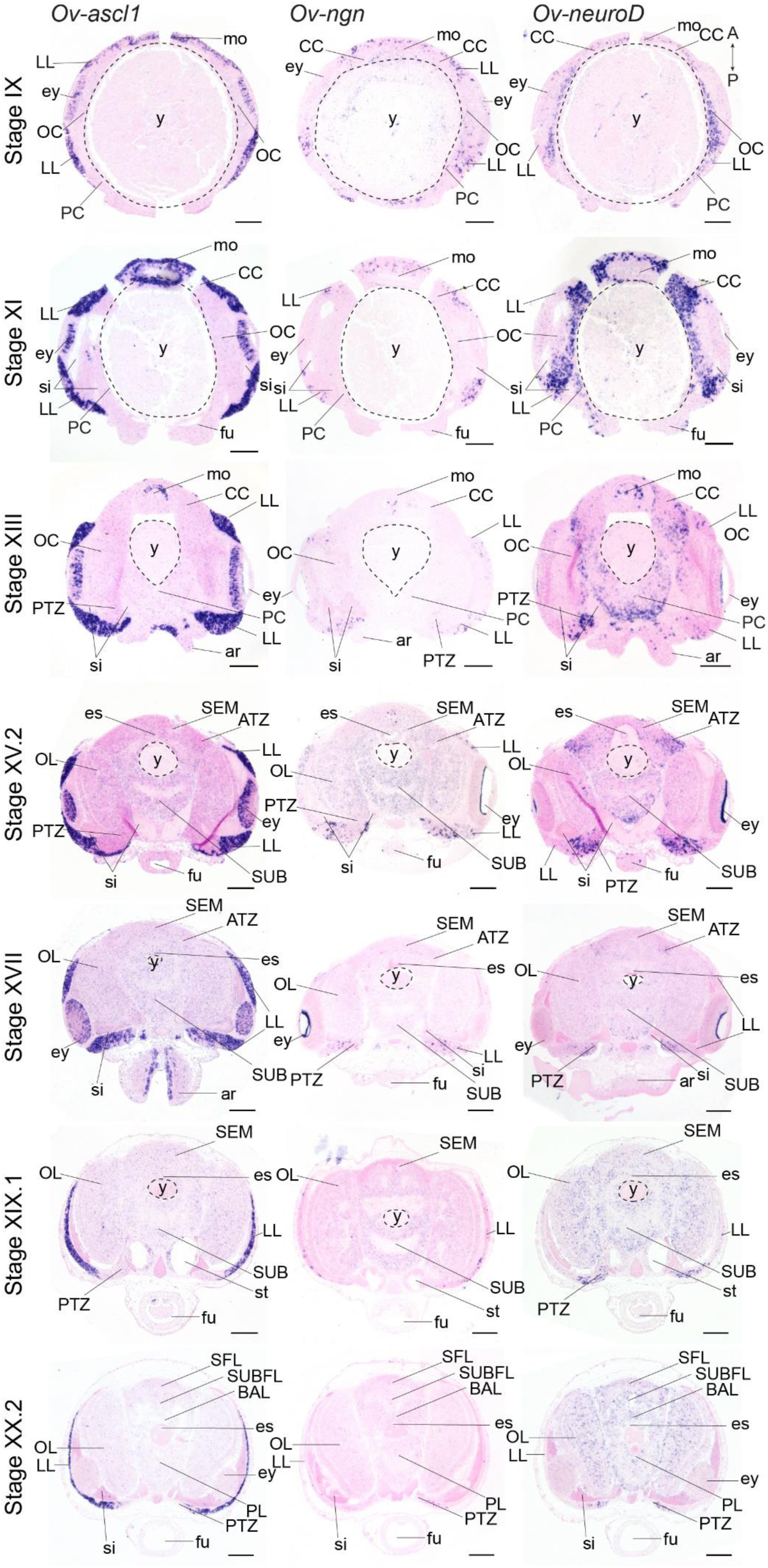
Expression of *O. vulgaris* proneuronal and neuronal differentiation bHLH genes during embryonic development. Expression of *Ov-ascl1* (left column), *Ov-ngn* (middle column) and *Ov-neuroD* (right column) on paraffin sections from Stage IX to Stage XX.2. *Ov-ascl1* is highly expressed in the lateral lips and retina at all stages. Overall, the number of *Ov-ascl1* positive cells increases during organogenesis, reaching a peak at Stage XV.2. Expression of *Ov-ngn* is restricted to cells in the lateral lips and a limited number of cells in the transition zones at all stages. *Ov-neuroD* is expressed at low level in the lateral lips at Stage IX and XI, but not in its most outer cell layers. At subsequent stages, *Ov-neuroD* is highly expressed in the transition zones. In the central brain, *Ov-neuroD* transcripts are present in the cerebral, optic and palliovisceral (FigS4.7) cords at Stage IX and also in the pedal cord from Stage XI onwards. Expression in the cords decreases over the course of development. Scale bars represent 100 μm. *Abbreviations as in* Figure 1.

The second atonal-related bHLH transcription factor studied here was NeuroD (Figure 4, S4.7). Its vertebrate and annelid homologs are expressed at the time neurons differentiate (Sur et al., 2017; Vervoort & Ledent, 2001). At the beginning of organogenesis (Stage IX and XI), *Ov-neuroD* was expressed in the cerebral, palliovisceral, pedal and optic cords of the central nervous system. Transcripts were also detected in the lateral lips, but at low level only. At Stage XI in the optic cords, a medio-lateral expression gradient was observed with high *Ov-neuroD* at the medial side and low *Ov-neuroD* at the lateral side, closer to the eyes. *Ov-neuroD* also seemed more highly expressed in the cerebral and palliovisceral cords compared to the pedal cord. At Stage XIII, *Ov-neuroD* transcripts were absent from the region of *Ov-ascl1* expressing cells in the lateral lips, but were present adjacent to those cells on the posterior side of the embryo next to the sinus ophthalmicus, and mark the posterior transition zone. In addition, *Ov-neuroD* expression in the cerebral, palliovisceral and optic cords was significantly reduced compared to earlier stages and expression in the pedal cord was elevated. At Stages XV.2 and XVII, high level *Ov-neuroD* expression marked both the anterior and posterior transition zones. In the central brain, transcripts were present at low level in the optic lobes, supra- and subesophageal masses. A similar expression pattern was visible at Stages XIX.1 and XX.2 with low level expression in all brain masses and clear presence in the transition zones. *Ov-neuroD* transcripts thus label the transition zones that connect the proliferative lateral lips to the postmitotic central brain.

While our data indicate that the lateral lips are a proliferating region with cells expressing the typical neurogenic transcription factors *Ov-ascl1* and *Ov-ngn,* it is not clear whether these represent different progenitor types. In addition, it was not proven yet that *Ov-neuroD* effectively labeled postmitotic cells. In a hybridization chain reaction experiment combined with immunohistochemistry (Figure 5), we show that *Ov-ngn* and *Ov-ascl1* are likely expressed in a different subset of progenitor cells, since their expression was not overlapping (Figure 5F). Furthermore, co-staining with the PH3 antibody demonstrated that *Ov-ascl1^+^* progenitor cells seem more proliferative compared to *Ov-ngn^+^* progenitors, that rarely overlap with the PH3^+^ population (Figure 5A-G). The proximity of *Ov-ngn^+^* cells to dividing *Ov-ascl1^+^* progenitors could be a sign of asymmetric progenitor division during neurogenesis in octopus. In addition, *Ov-neuroD* expressing cells did not co-localize with PH3 immunoreactive cells, indicating that *Ov-neuroD* is absent from mitotically active cells in octopus embryos (Figure 5H-M). If the progenitor cells in the lateral lips identified in this study would contribute neurons to the CNS, postmitotic neurons would need to travel long distances from the lateral lips to the central brain. However, direct proof of such neuronal cell migration is still lacking.

**Figure 5.**
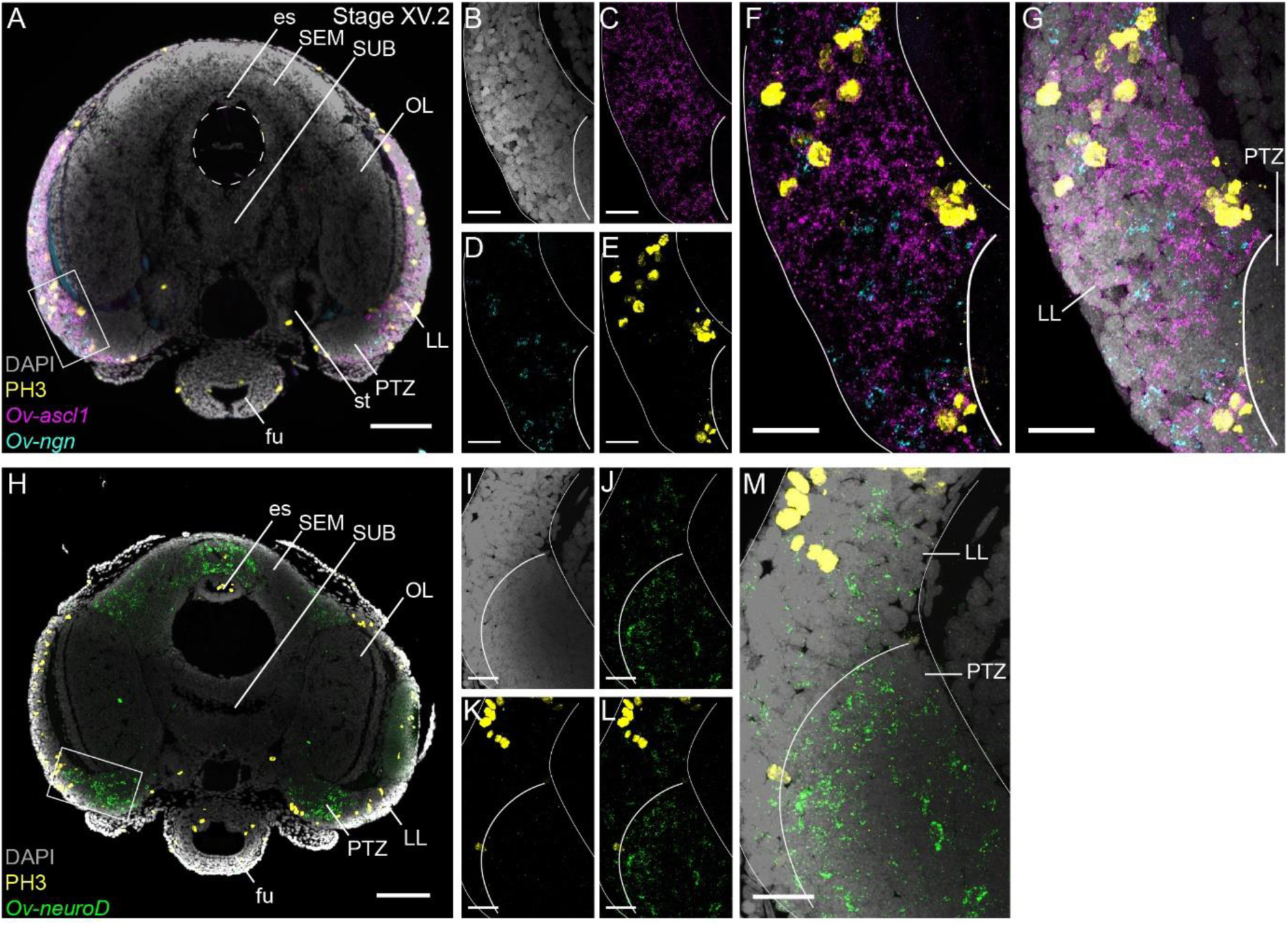
Cell proliferation profiles of the progenitor populations in the lateral lips. Multiplex *in situ* hybridization (HCR v3.0) combined with immunostaining against PH3. A. Overview image showing expression of *Ov-ascl1* and *Ov-ngn* and presence of mitotic cells (PH3+) on a transversal section of a Stage XV.2 embryo. The boxed area covering the lateral lips and posterior transition zone indicates the magnified region in B-G. B-G. Single- and multi-channel magnifications show that *Ov-ascl1* and *Ov-ngn* are expressed in different cell types. In addition, PH3 immunoreactivity is more common in *Ov-ascl1* expressing cells compared to *Ov-ngn* expressing cells. H. Overview image showing expression of *Ov-neuroD* and presence of mitotic cells on a transversal section of a Stage XV.2 embryo. The boxed area covering the lateral lips and posterior transition zone indicates the magnified region in I-M. I-M. Single- and multi-channel magnifications show that *Ov-neuroD* is broadly expressed in the PTZ and does not co-localize with dividing cells in the lateral lips. Scale bars represent 100 µm in A,H and 20 µm in B-G,I-M. *Abbreviations as in* Figure 1.

### Long distance neuronal migration from spatially patterned lateral lips to the developing brain

In order to map the progeny of cells generated in the lateral lips, we performed lineage tracing experiments using the fluorescent dye CFDA-SE. This technique has been applied *in vivo* to study temporal neurogenesis patterns in the mammalian cerebral cortex because of the short lifetime of the dye when not incorporated in cells and thus high temporal specificity (Govindan et al., 2018). We labeled different populations of cells in the lateral lips in early, mid or late organogenesis phases (Stages IX, XII.1 or XV.2) and traced their progeny to the maturation phase (Stage XIX.2) (Figure 6). First, we will focus on the optic lobe and peduncle complex (upper panels in Figure 6A,D,G). For these regions, we identified progenitor cells in the dorsal-anterior quadrant of the lateral lips that gave rise to cells in the optic lobes, in which labeled cells generally located to the inner and outer granular layers of the cortex (black populations with an asterisk in Figure 6; example of progeny in Figure 6E). Progenitor cells in the posterior lateral lips at Stages IX and XII.1 generated optic lobe cells that resided in the medulla, while at Stage XV.2, more cells located to the optic lobe cortex as well (example in Figure 6H). Ventral-anterior lateral lip progenitors did not generate optic lobe cells. We also identified a clear spatial patterning of progenitors that generate cells for the peduncle complex (olfactory & peduncle lobe). Progenitors in the posterior and ventral lateral lips generated cells destined to this complex (example in Figure 6H), while populations on the dorsal-anterior side did not.

**Figure 6.**
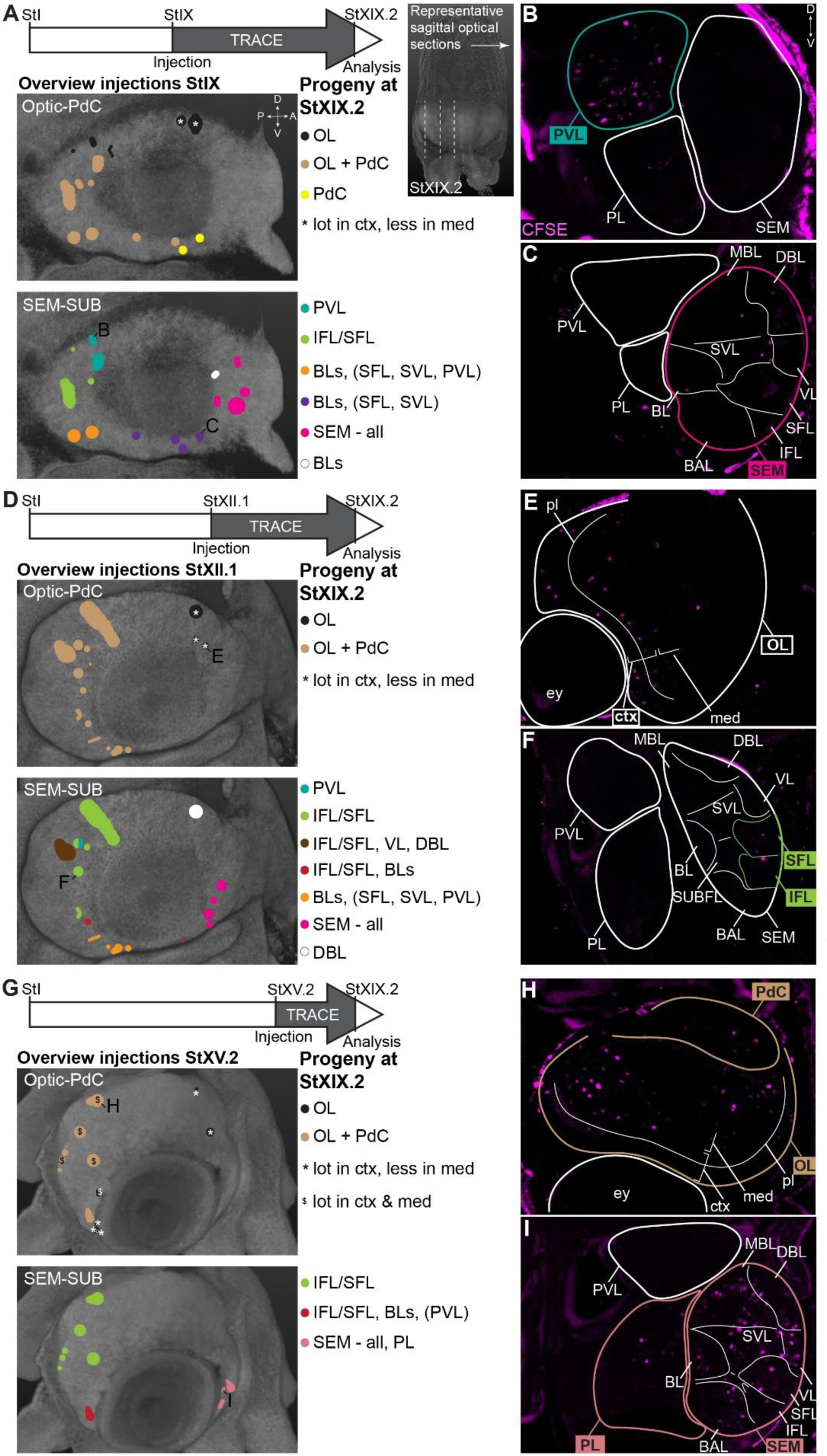
CFDA-SE lineage tracing from the lateral lips to the central brain. Injection of CFDA-SE in the lateral lips at Stage IX (A), Stage XII.1 (D) or Stage XV.2 (G) and tracing until Stage XIX.2 resulted in labeled cells in specific brain regions, depending on the location of the progenitor domain. Panels on the left show the location of the CFDA-SE injection site in the later lips, with each colored domain representing a single experimental condition. Progenitor cells in domains with the same color generated comparable output to the brain at Stage XIX.2. On the right, panels B,C,E,F,H,I display representative optical sections through the central brain, showing the differential output related to the position of the labeled progenitor population. Targeted regions are indicated in the color corresponding to the color-coded progenitor populations. *Abbreviations: BLs, basal lobes (BL, DBL, MBL); ctx, cortex; PdC, peduncle complex; others as in* Figure 1.

Considering output to the supra- and subesophageal masses (lower panels in Figure 6A,D,G; see Figure 1E-F for an overview of the lobes in the SEM and SUB), we observed that the posterior lateral lip progenitor cells specifically contributed cells to the inferior and superior frontal lobes of the supraesophageal mass at all stages, which has not been reported before by Koenig et al. (Koenig et al., 2016) (example in Figure 6F). Progenitor cells in the ventral lateral lips also produced cells destined to the basal lobes (basal lobe, dorsal basal lobe, medial basal lobe) of the supraesophageal mass (red and orange injections in Figure 6). The majority of cells located in the supraesophageal mass, however, were derived from ventral-anterior progenitor cells in the lateral lips (example in Figure 6I). In contrast to the supraesophageal mass and the optic lobe, few labeled progenitor populations generated cells for the subesophageal mass. A major contribution to the palliovisceral lobe found its origin in progenitor cells in the dorsal-posterior quadrant of the lateral lips at Stage IX, as was suggested but not empirically proven by Koenig et al. (Koenig et al., 2016) (example in Figure 6B). Other ventral-posterior progenitor populations generated a limited output to the palliovisceral lobe (orange and red injection spots in Figure 6). In addition, apart from the ventral-anterior progenitor populations at Stage XV.2 (example in Figure 6I), we did not identify progenitor populations that gave rise to a significant number of cells that migrated to the pedal lobe of the subesophageal mass. Occasionally, we identified few, single randomly dispersed cells in the pedal or palliovisceral lobes originating from more ventrally located progenitor populations. Considering the very low number of cells compared to the major output from those progenitors, these cells were not taken into account as they might have been labeled while migrating through another area. Taken together, our lineage tracing study identified spatial patterning in the lateral lips, that generate neurons for specific brain regions.

To determine the trajectory that the progeny of lateral lip cells is taking before entering the brain, we performed a short-term lineage tracing study. Hereto, populations of cells in the lateral lips were labeled with CFDA-SE at Stage XIV. Embryos were then allowed to grow for 48-72 hrs (reaching Stage XV.2) at which point they were fixed, cleared and imaged with a light sheet microscope to map the location of CFSE positive cells (Figure 7A, S7.1). We then manually tracked the labeled cells, and reconstructed their trajectory that revealed a continuous stream of cells starting in the lateral lips, passing the posterior (and dorsal) side of the lateral lips and the posterior transition zone, before entering the optic lobe (Figure 7B,C). On a series of optical sections, the trajectory can be followed in 2D (Figure 7D-R). Labeled cells in the lateral lips (intense labeling, Figure 7G-L) divided and migrated posteriorly and entered the posterior transition zone (labeling intensity decreased, Figure 7D-E). Then, cells could be traced towards the ventral side of the embryo in the posterior transition zone (Figure 7F-P) after which they occupied all layers in the optic lobe (Figure 7G-R). Populations labeled at a different location in the lateral lips showed similar trajectories (even the most anterior labelled population in Figure S7.1), with cells passing the dorsal and posterior lateral lips before entering the posterior transition zone and then the optic lobe. While this is clearly an important route, a (limited) trajectory passing through the anterior transition zone cannot yet be excluded. Taken together, cells destined for the optic lobe all seem to take a defined path via the posterior transition zone.

**Figure 7.**
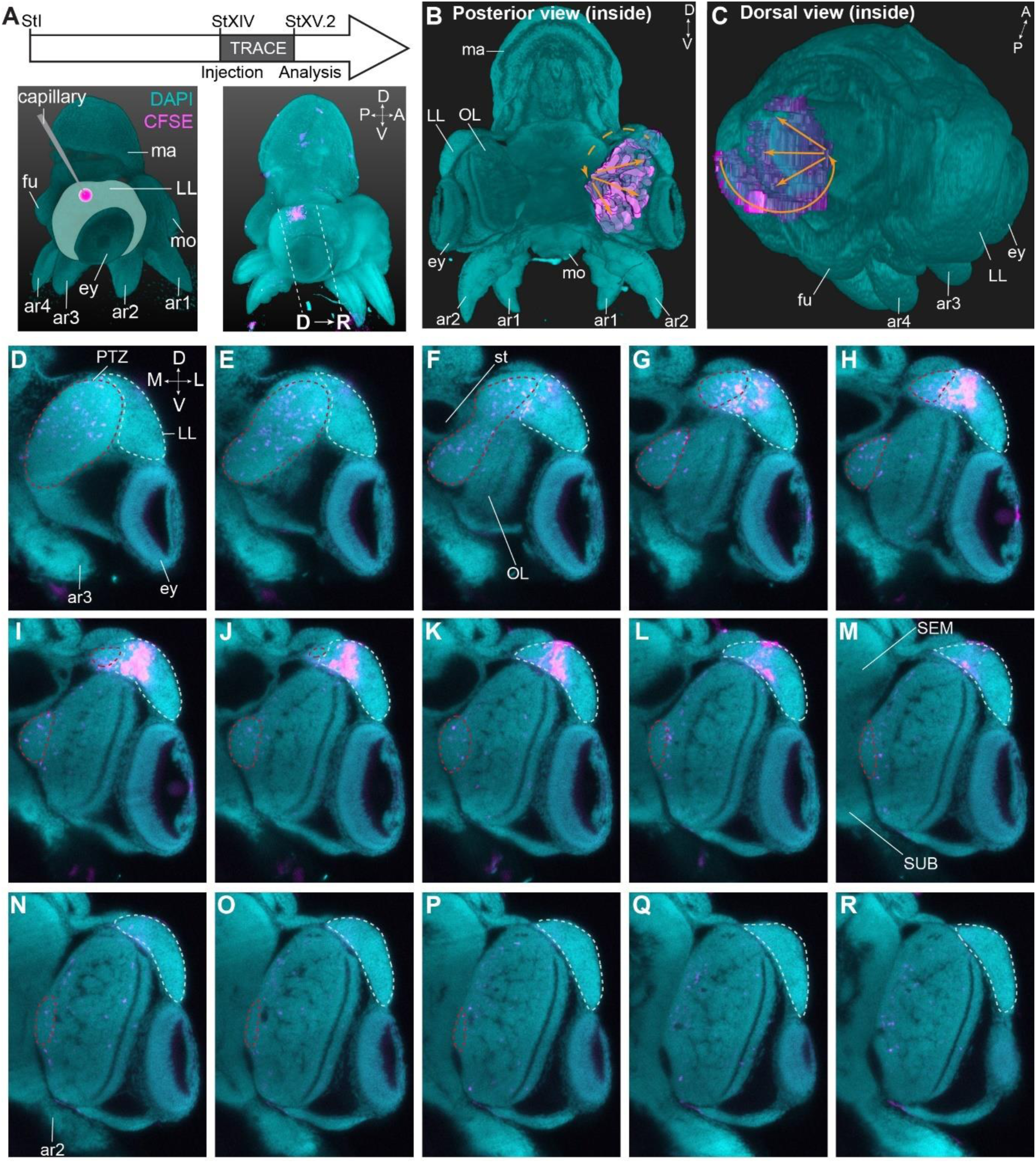
Trajectory mapping of migrating cells from the lateral lips to the central brain. A. Experimental setup showing CFDA-SE injection in the lateral lip at Stage XIV and embryo sampling at Stage XV.2. Dashed lines indicate optical sectioning planes for D-R. B-C. Volumetric rendering of the embryo (DAPI, cyan) and the manually traced CFSE object (pink-purple), showing the trajectory (orange arrows) from the injected area in the lateral lips to the optic lobe. D-R. Optical sections through the optic lobe from posterior (D) to anterior (R). The population of labeled cells in the lateral lips is visible in G-L. The progeny enters the optic lobes via the posterior transition zone. The lateral lips are encircled in mint green and the posterior transition zone in red. *Abbreviations as in* Figure 1.

## Discussion

This study showed that grossly, the embryonic octopus brain does not contain dividing progenitor cells. Instead, we identified a transient embryonic structure surrounding the developing eye – the lateral lips – that harbors proliferative cells expressing conserved pro-neural transcription factors. We further delineated embryonic neurogenesis in *O. vulgaris* using neural progenitor, pan-neuronal and differentiated-neuron marker genes with conserved functions in neurogenesis, neural specification and differentiation in protostomes and deuterostomes. The spatiotemporal expression patterns of these neurogenic genes suggest their involvement in regulating the development of the CNS in *O. vulgaris*. Koenig et al. have suggested a basic fate map of the neural primordia in *D. pealeii* using DiI tracing experiments, and identified contributions of cells from the lateral lips to the developing central brain (Koenig et al., 2016). Our data using a carboxyfluorescein ester that is less leaky compared to DiI, indicate that the spatial map seems to be conserved in cephalopods. In addition, we show that patterning is established at early neurogenesis stages and is temporally maintained, suggesting that lobe-specificity is determined early on within the progenitor area. Next to clear spatial patterning in the lateral lips, we identified long-distance migration of cells generated in the lateral lips towards the central brain in octopuses.

### *Octopus vulgaris* central nervous system maturation is delayed compared to other cephalopods

To determine when and where the first postmitotic immature neurons and differentiated neurons are formed in *O. vulgaris* embryos, we studied the expression of *Ov-elav* and *Ov-syt*, respectively. In the organogenesis phase, *Ov-elav* is expressed in embryonic cells lining the yolk envelope that form the cords, as in *O. bimaculoides* (Shigeno et al., 2015). The expression pattern is also consistent with the description of brain precursor regions proposed by Marquis after cytological studies in *O. vulgaris*, reinforcing the use of *elav* as a reliable marker for young octopus neurons (Marquis, 1989). We observed low level expression in all cords at Stage IX. In *O. bimaculoides*, expression was already reported from Stage VII.2 onwards, pointing towards rapid neuronal differentiation (Shigeno et al., 2015). In contrast to both octopus species, *Sof-elav1* expression in the cuttlefish *Sepia officinalis* is unequally distributed over the different cords, and disappears towards the end of embryonic development, pointing to a different timing of neuronal differentiation and a more advanced maturation of the *Sepia* brain at hatching (Buresi et al., 2013). Consistent with this, *S. officinalis* embryos have been described to respond to tactile and chemical as well as visual cues from within the egg capsule from Stage XV.1 and XVI onwards, respectively (Romagny et al., 2012). In our hands, *O. vulgaris* embryos only seem to react to visual stimuli (chromatophore contraction in response to light change) and mechanical stimuli (mantle contraction after tapping the chorion) from Stage XIX.1 onwards (preliminary observation). Furthermore, neural processes visualized with immunostaining against acetylated alpha-tubulin revealed the presence of neurites in the cerebral, palliovisceral, pedal and optic cords of *O. bimaculoides* and *S. officinalis* embryos as early as Stage VIII, while we only observed expression of *Ov-syt* and presence of acetylated alpha-tubulin in the central brain from Stages XIII and XV.2 onwards, respectively (Baratte & Bonnaud, 2009; Shigeno et al., 2015). These findings further support the delayed maturation of the brain in *O. vulgaris*, that perhaps uses its paralarval phase to complete maturation of the nervous system.

### Model of *O. vulgaris* neurogenesis and the specification of neural cell types

*Ov-soxB1* expression is not restricted to neurectodermal progenitor regions. Consistent with *Sof-soxB1* expression in *S. officinalis*, *Ov-soxB1* is expressed at high level in the surface ectoderm, in the developing eyes, and is absent from the gills and stellate ganglia (Focareta & Cole, 2016). Next to (neur)ectodermal expression, *soxB1* is also expressed in sensory epithelia in vertebrates, invertebrate mollusks and acoelomate worms (Focareta & Cole, 2016; Guo et al., 2010; Kiernan et al., 2005; Le Gouar et al., 2004; Neves et al., 2007; Semmler et al., 2010). In addition, SOXB1 proteins are present in both neurectodermal stem cells and differentiated neurons in certain species. *Drosophila soxN* and *Schmidtea polychroa soxB1* for example are expressed in early specification events in the CNS, but also in differentiated parts where they are involved in neuronal differentiation and axonal patterning, suggesting a dual role for protostome soxB1 (Ferrero et al., 2014; Girard et al., 2006; Monjo & Romero, 2015; Phochanukul & Russell, 2010). While such a general function in neuronal differentiation of vertebrate soxB1 factors has not been shown, some subtypes of (inter)neurons do require SOXB1 proteins for proper specification and migration (Cavallaro et al., 2008; Ekonomou et al., 2005; Panayi et al., 2010). Similar to many other species, our expression data suggest a dual role for *Ov-soxB1*, in early nervous system development to specify neural fate, and later on to steer neural cell differentiation.

After neurectoderm establishment and neural stem cell formation regulated by SOXB1 transcription factors, neural progenitors must be specified. In most bilaterians, this function has been attributed to members of the bHLH family of transcription factors, including *atonal*, *neurogenin*, *neuroD* and *achaete-scute* subfamilies. Our study is the first one mapping the expression of *ascl1* in cephalopods and together with the expression pattern of *Ov-ngn*, suggests the presence of a neurogenic domain in the lateral lips, outside the cords of the central brain (Figure 8A,B). In the annelids *Capitella teleta* and *dumerilii*, *neurogenin* and *ash1* are both expressed in actively proliferating neural progenitor cells in the neurectoderm (Figure 8C). While *Ct-ngn* positive cells stay on the apical side, the *Ct-ash1* positive progenitors ingress and undergo limited division before becoming *Ct-elav* positive (Demilly et al., 2013; Meyer & Seaver, 2009; Simionato et al., 2008; Sur et al., 2017, 2020). In *O. vulgaris*, the *Ov-ascl1* progenitor population seems to be more proliferative compared to the *Ov-ngn* progenitor population, which suggests their sequential expression could be turned around in cephalopods (Figure 8B,C). In addition, the *Ov-elav* expressing neurons in the cords are not located immediately basal from the *Ov-ascl1* or *Ov-ngn* positive pool of neural progenitor cells. In particular, a population of *Ov-neuroD* expressing cells is found in the transition zones, in between the *Ov-ascl1^+^* and *Ov-ngn^+^* lateral lips and the developing *Ov-elav^+^* brain, suggesting an intermediate, *Ov-neuroD^+^* population is present in *O. vulgaris* (Figure 8A,B). While lost in *Drosophila* and *Ciona intestinalis*, *neuroD* genes can be widely found in bilaterians, where they steer the differentiation of neurons (Ledent et al., 2002; Stollewerk & Simpson, 2005). In contrast, in the trunk of the annelid *P. dumerilii* (but not *C. teleta*) and in the developing nervous system of the planaria *S. polychroa* and *S. mediterranea*, *neuroD* seems broadly expressed in the neurectoderm, which suggests that the role of neuroD might not be conserved in Spiralia (Figure 8C) (Cowles et al., 2013; Meyer & Seaver, 2009; Monjo & Romero, 2015; Simionato et al., 2008; Sur et al., 2020). However, co-localization studies of *neuroD* with progenitor marker genes are still lacking in these species. In vertebrates, *ascl1* and *ngn* are expressed in a complementary fashion, both exerting proneuronal functions in neurogenesis (Bertrand et al., 2002; Castro et al., 2011; Farah et al., 2000; Gradwohl et al., 1996; Lee et al., 1995; Ma et al., 1996). Similar to our data in octopus, n*eurogenin* expression precedes, but also partially overlaps with that of *neuroD* (Grimaldi et al., 2008). In contrast, *Drosophila* has only one *neurogenin/neuroD* homolog called *tap*, which relates better to *neurogenin* based on sequence similarities, but does not have a proneural role and is expressed in a few neurons, regulating their differentiation, axonal growth and guidance (Gautier et al., 1997; Vervoort & Ledent, 2001; Yuan et al., 2016). In arthropods, the members of the *achaete-scute* subfamily are expressed in proneural clusters and promote the generation of neural progenitor cells from quiescent ectodermal cells (Bertrand et al., 2002; Cubas et al., 1991; Quan & Hassan, 2005; Skeath & Carroll, 1991; Stollewerk & Chipman, 2006). In cnidaria as well, *ashA* promotes neurogenesis during its development (Layden et al., 2012) (Figure 8C). In *O. bimaculoides*, *neurogenin* and *neuroD* transcripts were detected in the prospective cerebral, palliovisceral and pedal cords at Stage VIII, based on whole mount *in situ* hybridization, but did not distinguish the lateral lips from the cordal areas (Shigeno et al., 2015). Based on cross sections, *neurogenin* transcripts were found on the surface of the embryo, and *neuroD* more at the level of the cords, similar to our observations in *O. vulgaris* (Shigeno et al., 2015). In the vertebrate neural tube, NEUROD was also found on the basal side, in postmitotic neuronal and glial cells and is required for the differentiation of neurons in the inner ear, cerebellum and hippocampus (reviewed in Dennis, Han and Schuurmans, 2019). Its expression pattern in *O. vulgaris* suggests that upon differentiation, neurons express *Ov-neuroD* before expressing *Ov-elav*, which is similar to the role of *Ov-neuroD* as neuronal-differentiation bHLH transcription factor, as described for vertebrates and the annelid *C. teleta* (Bertrand et al., 2002; Farah et al., 2000; Lee et al., 1995; Sur et al., 2020). We also revealed possible overlapping expression of *Ov-soxB1* with *Ov-ascl1* and *Ov-ngn* in the lateral lips throughout development, suggesting a similar activation of *Ov-ascl1* and *Ov-ngn* by *Ov-soxB1* in the presumptive neurectoderm (Figure 8B) (Amador-Arjona et al., 2015; Ferrero et al., 2014). Therefore, *Ov-ngn* and *Ov-ascl1* are expressed at the right time and place to be the major proneuronal genes for the formation of the central nervous system in *O. vulgaris*. Our data further substantiate the conserved expression of proneural bHLH transcription factors, which suggests they might already have been present in the ur-bilateria.

**Figure 8.**
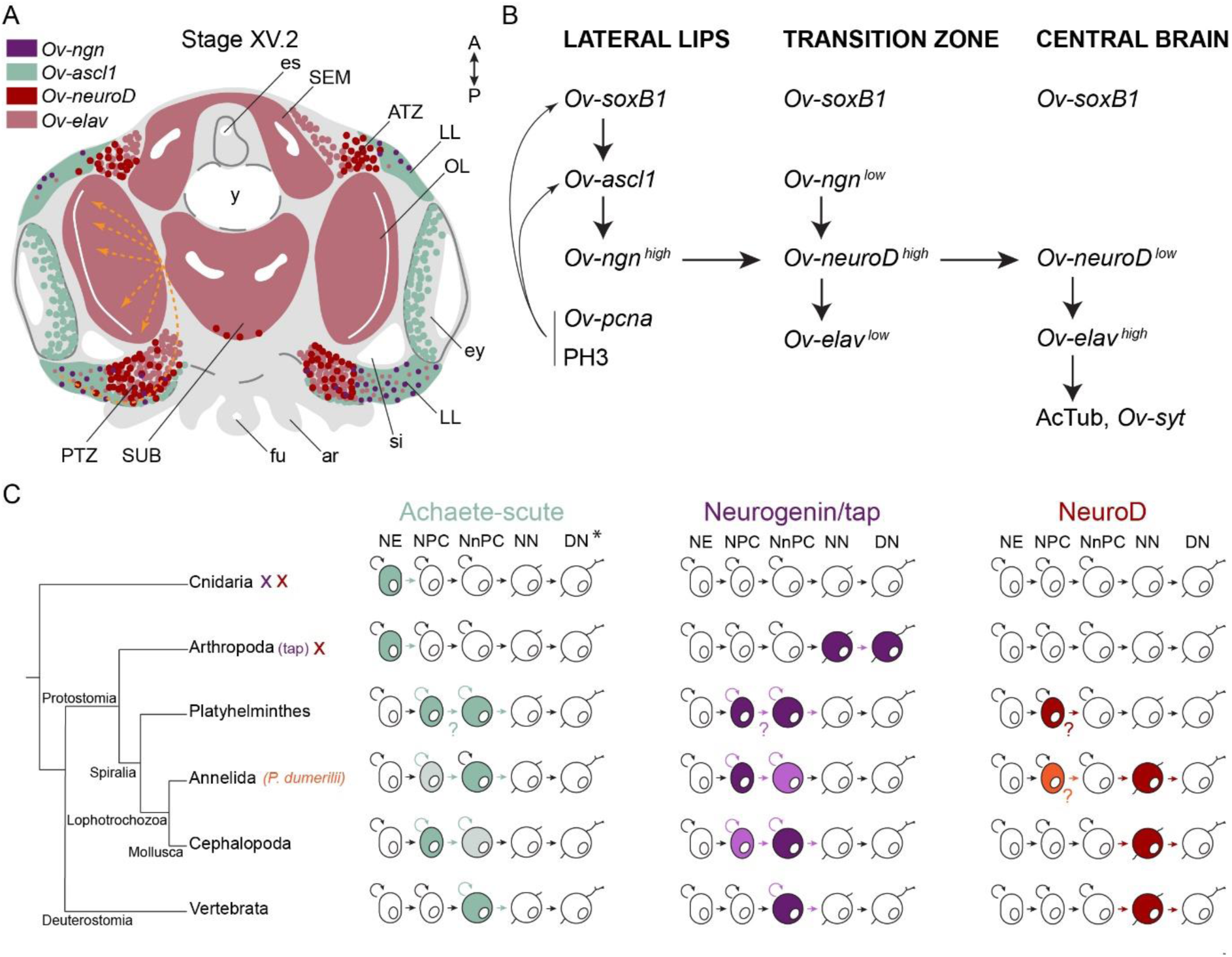
Overview of the expression of neurogenic genes and hypothetical neurogenesis process in *O. vulgaris*. A. The expression domains of *Ov-ngn* (purple), *Ov-ascl1* (green), *Ov-neuroD* (red) and *Ov-elav* (pink) are depicted during *O. vulgaris* neurogenesis. Areas indicate high level expression in most cells whereas dots represent lower expression or high expression in a couple of cells. The orange arrows depict the trajectory taken by cells originating in the lateral lips, passing through the posterior transition zone before entering the optic lobe, as observed after CFDA-SE lineage tracing. The arrows are dashed, considering their 3D projection on a 2D figure. B. *Ov-soxB1*, *Ov-ngn* and *Ov-ascl1* are all expressed at high level in the lateral lips. In this structure as well, proliferating cells (*Ov-pcna* expressing or PH3 positive) are abundant. They mostly colocalize with *Ov-ascl1* expressing cells that potentially also express *Ov-soxB1*. Our data suggest the onset of differentiation in the transition zones, with low-level expression of *Ov-ngn*, high-level expression of *Ov-neuroD* and, low-level expression of *Ov-elav*. We suggest that the induction of *Ov-neuroD* expression is guided by *Ov-ngn* and that differentiating cells express *Ov-neuroD* before *Ov-elav*. Once arrived in the central brain, neurons start forming synapses (*Ov-syt* expression, presence of acetylated alpha-tubulin). C. Evolutionary comparative expression mapping based on a generalized neural specification and differentiation sequence. See main text for details and references. *Note that the depicted cell types are a generalized state, and that certain phyla/subphyla/classes might lack one or more progenitor types. *Abbreviations: ar, arm; ATZ, anterior transition zone; DN, differentiated neuron; es, esophagus; ey, eye; fu, funnel; LL, lateral lips; NE, neurectodermal cell; NN, newborn neuron; NnPC, neuronal progenitor cell; NPC, neural progenitor cell; OL, optic lobe; PTZ, posterior transition zone; SEM, supraesophageal mass; si, sinus ophthalmicus; SUB, subesophageal mass; y, yolk*.

### Spatial patterning of the lateral lips and long-distance migration to the central brain

Proliferating cells with a signature of neural progenitors are thus located in the lateral lips, while more mature neurons are found in the cords. The absence of proliferative cells in the developing brain in an organism with such a big and centralized CNS is striking, but not uncommon for mollusks. Mitosis in the *Aplysia* CNS is infrequent from early embryogenesis to adulthood (Jacob, 1984). Compared to other invertebrates with large brains such as insects, however, this seems to be a rather unique strategy. The developing optic lobe in *Drosophila* for example has dividing neuroblasts in two main proliferation centers inside the lobe. These neuroblasts had previously invaginated from the neuroepithelium and start dividing only when in the brain (Álvarez & Díaz-Benjumea, 2018; Apitz & Salecker, 2015; Green et al., 1993; Hofbauer & Campos-Ortega, 1990; Walsh & Doe, 2017). The physical disconnection between the neural stem cells and their progeny in octopus suggests that secondary progenitor cells or neurons migrate towards the central brain where they integrate.

In lophotrochozoans such as the annelids *C. teleta* and *P. dumerilii,* dividing neural progenitors are located on the apical side of the neuroepithelium and it is their progeny that ingresses to form the CNS (Meyer & Seaver, 2009; Monjo & Romero, 2015; Sur et al., 2020). While resembling the postmitotic migration in *O. vulgaris*, their migratory path is short and the number of neurons remains limited, making direct comparison with *O. vulgaris* difficult. As described here and by Shigeno et al., the nervous system of cephalopods originates from a system of cords, similar to the nervous system of primitive mollusks like Aculifera including chitons, and in contrast to Concifera including gastropods, whose CNS knows a ganglionic origin (Richter et al., 2010; Shigeno et al., 2015; Sumner-Rooney & Sigwart, 2018). Gastropod ganglia seem to derive from ectodermal cells in the body wall that proliferate and delaminate/migrate inwards to join the developing ganglia. These studies suggest that similar to annelids, neural progenitor cells in the gastropod mollusk *Aplysia californica*, divide in proliferative zones in the body wall and their progeny migrates few cell lengths to the nearest ganglion (Demian & Yousif, 1975; Jacob, 1984). Depending on the location in the proliferative zone, cells migrate either in a columnar stream, or individually using pseudopodia. Evidence for cell migration in cephalopods came from *Loligo vulgaris*, where *in vitro* cultures of the “oculo-ganglionar complex” showed extensive migration of cells and differentiation into bi-and multipolar neurons (Marthy & Aroles, 1987). Here, we showed that the progeny of octopus lateral lip progenitors migrates over long distances before integration in the CNS. Strikingly, we observed that the migratory path does not just represent the shortest route to destiny. Instead, we found that independent from the anterior-posterior axis, progenitor populations in the dorsal lateral lips generate cells for the optic lobe that pass through the posterior transition zone. Cells thus migrate from the most anterior part towards the posterior part of the lateral lips before entering the posterior transition zone and eventually migrating in all directions, spreading throughout the optic lobe. This suggests active, directed migration controlled by guidance cues. Which cell intrinsic and/or extrinsic cues govern the migratory process, remains to be studied.

Active neural migration guided by extrinsic cues is common in vertebrates that build their brain from a large, spatially patterned and proliferating neural epithelium folded into a tube. Neurons are generated in a temporally controlled fashion and migrate away from the progenitor zone surrounding the ventricles to form the grey matter. The latter entirely consists of postmitotic neurons that start growing neurites and connect while additional postmitotic neurons are migrating in between them to their target region (Marín et al., 2010; Marín & Rubenstein, 2003; Paridaen & Huttner, 2014; Taverna et al., 2014). In that respect, the development of the *O. vulgaris* brain seems strikingly similar. Yet *O. vulgaris* seems to pattern the lateral lips in such a manner that entire brain lobes are derived from specific areas in the lateral lips, whereas specification in vertebrates links spatial (i.e. progenitor area) or temporal patterning to cell types that are intermingled in certain regions. An example is the subpallium, that generates (among other cell types) cortical interneurons, while the dorsal pallium generates pyramidal neurons, and both are mixed in the cerebral cortex. Another example is the postnatal V-SVZ, where different areas generate distinct interneuron types destined for the olfactory lobe (Hatten, 1999; Marín & Rubenstein, 2001, 2003; Medina & Abellán, 2009).

Neurons that are born in a specific region of the lateral lips thus seem instructed to migrate to a certain brain lobe, but how they get specified to different cell types within that lobe remains unclear. Indeed, cells from one injection site mostly spread out over a whole lobe, except for the optic lobe, where we identified lateral lip subregions that are biased to generate optic lobe cortex or optic lobe medulla neurons. Interestingly, the proportion of cells in those layers increased when progenitor labeling was performed at later stages, indicating that cell types might be specified in a temporal manner as well. To test this hypothesis, we first need to identify molecular markers for different neuronal subpopulations, as these have not yet been identified. Furthermore, while proneural bHLH transcription factors are likely involved in specifying and maintaining neural progenitor identity, the transcription factors that serve as terminal selectors need to be revealed.

## Materials and Methods

### Animals

Live *O. vulgaris* embryos were obtained from the lab of E. Almansa (IEO, Tenerife), transferred to the lab of Developmental neurobiology and kept in a closed standalone system (Deryckere et al., 2020). Embryos were observed, staged and sampled daily, followed by overnight fixation in 4 % paraformaldehyde (PFA) in phosphate buffered saline (PBS). After a wash in PBS, embryos were manually dechorionated with forceps and transferred to embedding cassettes (Tissue-Tek Biopsy 6-Chamber Cassette, Sakura). For paraffin processing, the cassettes were immersed in 0.9 % NaCl overnight before progressive dehydration and paraffin-embedding using an Excelsior^TM^ AS Tissue Processor and HistoStar^TM^ Embedding Workstation (Thermo Scientific). 6 μm-thick transversal sections were made for subsequent immunohistochemistry or *in situ* hybridization.

### Immunohistochemistry on paraffin sections

Embryo sections were processed using an automated platform (Ventana Discovery, Roche) for direct fluorescent staining. Primary antibodies mouse anti-Acetylated alpha Tubulin (Sigma T6793) and rabbit anti phospho-histone H3 (Ser10) (Millipore 06-570) and secondary antibodies donkey anti-mouse Alexa 488 and donkey anti-rabbit Alexa 555 (Life Technologies) were each diluted in antibody diluent (Roche) and incubated at a final concentration of 1:300. Sections were then incubated in DAPI and mounted in Mowiol. Images were acquired with a Leica DM6 upright microscope and minimum/maximum displayed pixel values were adjusted in Fiji (Schindelin et al., 2012).

### RNA extraction, sequencing and Iso-Seq data analysis

In order to construct a full-length transcriptome of *O. vulgaris* embryos and paralarval brains, the Iso-Seq method was used. RNA was extracted from a pool of 25 Stage XI-XII embryos using Tri-reagent (Invitrogen) and the Qiagen Micro kit (Qiagen). cDNA was synthesized with the Clontech SMARTer PCR cDNA Synthesis Kit (Takara Bio Inc). RNA was also extracted from dissected brains of one-day old paralarvae in a similar manner and cDNA was synthesized using the NEBNext cDNA Synthesis & amplification kit. Both samples were sequenced on the PacBio Sequel at the Genomics Core at KU Leuven (Belgium) following the protocol recommended by PacBio. Only cDNAs containing polyA-tails were selected, with the aim to retrieve full-length transcripts. This resulted in a total of 12,017,703 subreads for the embryo and 15,426,835 subreads for the brain sample. The raw data files were processed with SMRT Link release 9.0.0 software. The IsoSeq 3.3 pipeline was followed to generate consensus reads (inc. polish, min.passes =1). Lima (-isoseq) was used to retain full-length fragments that possess both primers only, to remove unwanted primer combinations and to orient the sequences. Subsequently, Poly(A) tails were trimmed and concatemers were removed. This resulted in 22,757 and 28,490 high-quality polished isoforms for the embryo and brain samples, respectively. Data have been deposited in SRA under the following accession number PRJNA718058.

### Identification and cloning of *O. vulgaris* genes

Putative homologs of *O. vulgaris achaete-scute*, *neurogenin*, *neuroD*, *elav*, *soxB1, synaptotagmin* and *pcna* genes were identified using tBLASTn searches against the ISOseq transcriptomes. *O. vulgaris* hits hereafter named *Ov-ascl1*, *Ov-ngn*, *Ov-neuroD*, *Ov-elav, Ov-soxB1*, *Ov-syt* and *Ov-pcna* were then blasted against the NCBI database to verify sequence homology. Primers were designed (primer sequences in Table S1) to isolate a 500-1000 bp fragment from mixed-stage *O. vulgaris* embryo cDNA (synthesized using Superscript III Reverse Transcriptase (Invitrogen)) (probe sequences in Table S2). The resulting PCR products were TA cloned into the pCR^TM^II-TOPO vector (Invitrogen) and sequenced by LGC Genomics (Berlin). After plasmid linearization, anti-sense digoxigenin-(DIG) labeled RNA probes were generated using an Sp6- or T7-RNA polymerase and DIG RNA labeling mix (both Roche) following the manufacturer’s protocol. The probes were cleaned using Micro Bio-Spin^TM^ P-30 Gel Columns with RNase-free Tris Buffer (BioRad).

### Colorimetric in situ hybridization

Paraffin sections were processed using an automated platform (Ventana Discovery, Roche) with RiboMap fixation and BlueMap detection kits (Roche) for *in situ* hybridization. In short, sections are deparaffinated, heated to 37 °C, post-fixed and pretreated. Then, a 4-minute digestion with proteinase K (Roche, 1:1000 in PBS-DEPC) is followed by probe titration (100-300 ng per slide dependent on the probe, dissolved in Ribohybe reagent (Roche)), denaturation at 90 °C for 6 minutes and hybridization at 70 °C for 6 hrs. Three stringency washes in 0.1X SSC at 68 °C for 12 minutes each are followed by post-fixation. The anti-DIG-Alkaline phosphatase antibody (Roche) is added and sections are incubated for 30 minutes after which a colorimetric signal (BCIP/NBT) is developed for 4-9 hrs (probe dependent). The tissue is counterstained with Red Counterstain II (Roche), followed by dehydration and mounting using Eukitt quick-hardening mounting medium (Sigma). Bright-field images were taken with a Leica DM6 upright microscope and background was subtracted in Photoshop. Images used in the figures represent the expression pattern observed in multiple embryos.

### Hybridization Chain Reaction v3.0

HCR-3.0-style probe pairs for fluorescent *in situ* mRNA visualization were generated for *Ov-ascl1*, *Ov-neuroD and Ov-ngn*. Hereto, the insitu_probe_generator (Null & Özpolat, 2020), followed by BLAST searches using Blast2GO (Conesa et al., 2005) to minimize potential off-target hybridization. DNA oPools were ordered from Integrated DNA Technologies, Inc. (probe sets in Table S3) and dissolved in DNase/RNase-Free distilled water (Invitrogen). HCR amplifiers with fluorophores B1-Alexa Fluor-546, B2-Alexa Fluor-647 and B3-Alexa Fluor-488 were ordered from Molecular Instruments, Inc. The Molecular Instruments HCR v3.0 protocol for FFPE human tissue sections, based on Choi et al. 2016 and 2018 was followed (Choi et al., 2016, 2018). Described here are adaptations from this protocol. Paraffin sections were baked at 65 °C for 30 min and subsequently deparaffinized with Xylene (2 x 4 min) and 100% EtOH (3 x 4 min). To permeabilize the tissue, slides were treated with proteinase K (Roche, 1:3000 in PBS-DEPC) for 5 min at 37 °C. Slides were then rinsed 2 x 2 min with autoclaved MQ and immediately processed for HCR. After a 30 min pre-hybridization step, probe solution (0.4 pmol per probe in probe hybridization buffer) was incubated overnight. The next day, 4.5 pmol of hairpin h1 and 4.5 pmol of hairpin h2 were snap-cooled (95 °C for 90 seconds, 5 minutes on ice followed by 30 min at room temperature) and added to 75 µL of amplification buffer. After overnight amplification, excess hairpins were removed by washing 3 x 10 min with 5X SSCT. After HCR, we proceeded with immunohistochemistry as described above and lastly, sections were incubated in DAPI and mounted in Mowiol. Images were acquired using a confocal microscope (Fluoview FV1000, Olympus) or an upright microscope (DM6, Leica) and minimum/maximum displayed pixel values were adjusted in Fiji (Schindelin et al., 2012).

### Phylogenetic analysis

To determine homology, phylogenetic analyses of ASH, NEUROD, NEUROG, ELAV and SOX families were performed. Full length protein sequences (when available) were obtained using BLASTp, tBLASTn or word search in NCBI or from published articles (accession numbers in Table S4). In the case of *E. scolopes*, the BLAST genome server together with peptide sequences on cephalopodresearch.org were used (Belcaid et al., 2019). In the case of *O. vulgaris*, nucleotide sequences were obtained using tBLASTn searches against the Iso-Seq transcriptomes, followed by a search for open reading frames using the ORFfinder tool in NCBI, which also provided the translated protein sequences (*O. vulgaris* protein sequences in Table S5). All protein sequences were aligned using the ‘MUSCLE Alignment’ feature (Edgar & Sjölander, 2004) within MEGA-X (Kumar et al., 2018). The matrix was then trimmed using TrimAI (Capella-Gutiérrez et al., 2009) via the *Automated 1* algorithm. The best-fit substitution model for each alignment was determined using the Bayesian information criterion in IQ-TREE (Nguyen et al., 2015). Maximum likelihood analyses using the LG+G4 (ELAV), JTT+I+G4 (NEUROD/NGN), VT+I+G4 (ASH) or JTT+G4+F (SOX) substitution models for protein evolution were also performed in IQ-TREE, with branch supports calculated by 10.000 Ultrafast bootstrap replicates (UFBoot2) from maximum 1000 iterations (stopping rule) (Hoang et al., 2018; Nguyen et al., 2015). The produced consensus trees were rooted and visualized with FigTree v1.4.4 (Rambaut, 2018). Domains in the predicted protein sequences of candidate *O. vulgaris* homologs were identified with NCBIs’ conserved domain database (CDD) search tool (Lu et al., 2020).

### CFDA-SE injection procedure

For injection of live *O. vulgaris* embryos, a glass capillary (3-000-203-G/X, Drummond), that was pulled using a Laser-Based Micropipette Puller (P-2000, Sutter Instrument; Heat 450, Fil 4, Vel 150), and opened at 30 μm was mounted on a micromanipulator (M3301L, WPI) and connected to a FemtoJet (Eppendorf) via an injection tube. Excess seawater was removed from the egg using a tissue after which the egg was transferred to a Sylgard-coated Petri dish (Sylgard 170, Dowsil). The egg was stabilized with tweezers and the capillary was inserted in the embryonic tissue, from the stalk side through the chorion, in an angle of about 10-30 ° relative to the dorso-ventral axis of the embryo. A single dose of 50 - 100 nL CFDA-SE working solution (1 mM for trajectory mapping, 0.1 mM for long term tracing in filtered sea water, 3% FastGreen) was injected in the lateral lips anterior, posterior, dorsal or ventral from the eye placode at developmental Stage XIV for trajectory mapping and Stages IX, XII.1 and XV.2 for long term tracing. After injection, embryos were placed back in a Petri dish with filtered sea water. Successful uptake of the dye was verified after 30 minutes using a fluorescence binocular (SteREO Discovery.V8 with AxioCam MRc5 (Zeiss)). Individual eggs were then incubated in a 96-well plate in filtered sea water, 2% Penicillin/Streptomycin, in a 19 °C incubator in the dark (Heratherm IMC18, Thermo Scientific). Viable embryos were sampled after 48 or 72 hrs for trajectory mapping or when reaching Stage XIX.2 for long term tracing, fixed overnight in 4 % PFA in PBS and then stored in PBS until dechorionation and clearing.

### Clearing and whole mount immunohistochemistry

Dechorionated embryos were cleared before light sheet imaging as previously described (Deryckere et al., 2020). Whole mount immunohistochemistry was performed for long term CFDA tracing, during the PBS washing steps in between incubation in Sc*ale*CUBIC-1 and Sc*ale*CUBIC-2 as follows: after clearing in Sc*ale*CUBIC-1, embryos were washed 4 times; 1 time for 5 mins and 3 times for 2 hours in PBS supplemented with 0.3% Triton X-100 (PBS-T) and then incubated in primary antibody solution in PBS-T (goat anti-fluorescein (Novus Biologicals NB600-493, 1:500, pre-incubated overnight with non-injected embryos)) for 2 days at 4 °C. Embryos were then washed 3 times for 2 hours in PBS-T, secondary antibody solution was added (donkey anti-goat Alexa 488 (Life Technologies, 1:300)) and incubated overnight at 4 °C after which the samples were washed 3 times for 2 hours in PBS-T and then incubated in 1/2-water diluted Sc*ale*CUBIC-2 at 37 °C.

### Light sheet fluorescence microscopy and analysis

Cleared and stained embryos were glued with their yolk sack on a metal plunger and imaged using a Zeiss Z1 light sheet microscope (Carl Zeiss AG, Germany) in low-viscosity immersion oil mix (Mineral oil, Sigma M8410 and Silicon oil, Sigma 378488, 1:1). Then, 3D reconstructions were generated in Arivis (Vision4D, Zeiss Edition 3.1.4).

For lineage tracing after CFDA-SE injection, the distribution of the progeny was mapped using optical sections. In addition, for trajectory mapping, the region including most CFSE labeled cells was manually traced using the objects drawing tool in Arivis, in order to visualize the trajectory in 3D. This same tool was also used to reconstruct the eye, optic lobe, lateral lips and posterior transition zone at Stage XV.2, and to reconstruct the different lobes in the central brain at hatching for Figure 1.

## Acknowledgements

The authors wish to thank Eduardo Almansa (Instituto Español de Oceanografía, Santa Cruz de Tenerife, Spain) and Camino Gestal (Institute of Marine Research, Vigo, Spain) for generously providing *Octopus vulgaris* embryos. We thank the laboratory of Cris Niell (Institute for Neuroscience, University of Oregon, USA) for sharing their Hybridization Chain Reaction protocol. We are also grateful to the Genomics Core Leuven (www.genomicscore.be) for the IsoSeq sequencing and Joke Allemeersch for data analysis. We wish to thank Maria Antonietta Tosches for critically reading the manuscript.

## Supplementary figures

**Figure S2.1.**
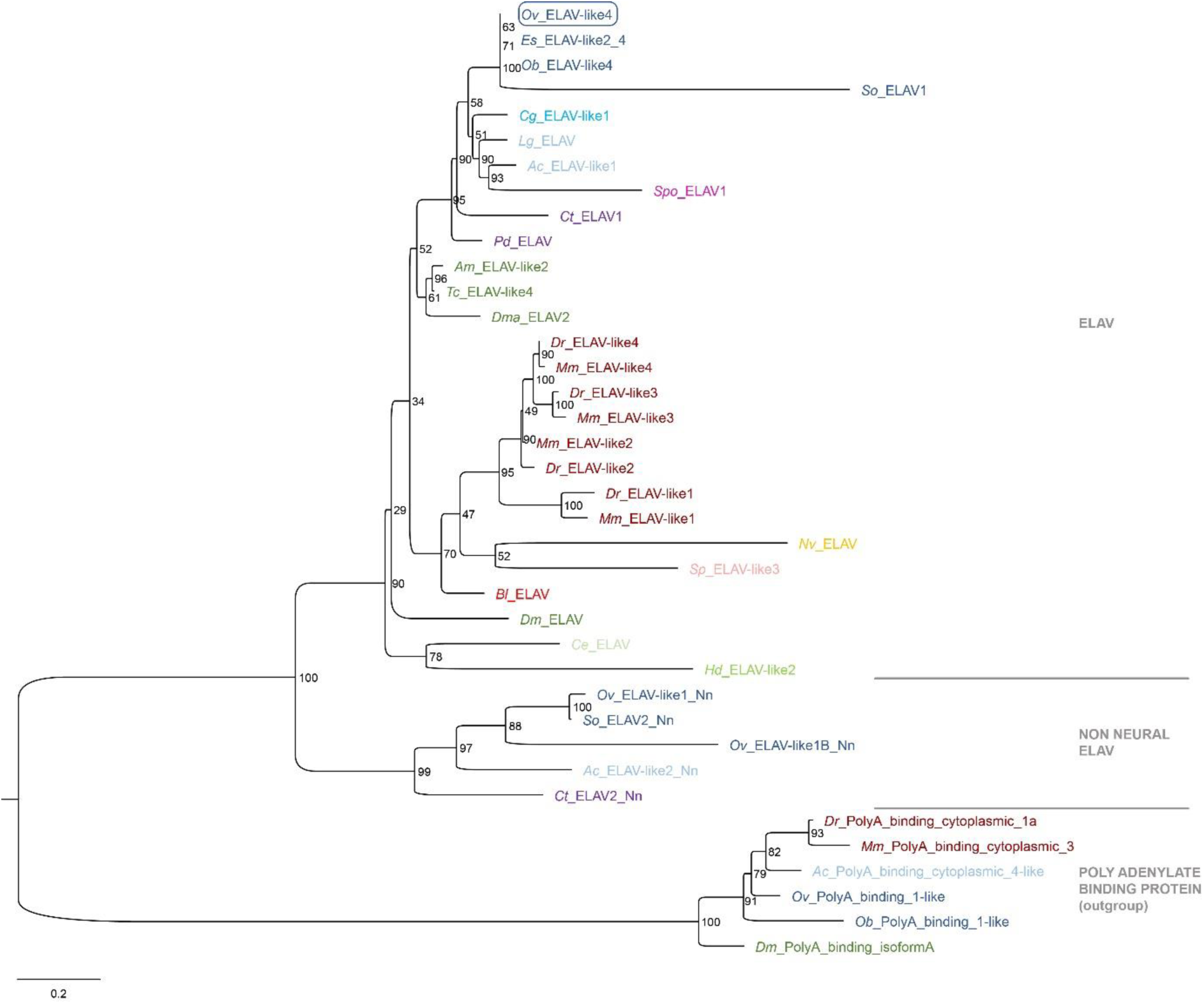
Phylogeny *Ov*-ELAV. Phylogenetic analysis of ELAV and ELAV-like proteins shown in a maximum likelihood tree. Numbers at each branch represent bootstrap support values. The tree has been rooted using the RNA binding protein Poly(A)-binding protein as outgroup. While vertebrates have four (or more) neural *ELAV* genes, invertebrates have only a single one. Lophotrochozoa and cnidaria, but not deuterostomia possess additional non-neural *ELAV*-like genes (Nn). The *O. vulgaris* ID used for subsequent ISH probe design is framed in blue. The scale bar represents the number of amino acid substitutes per site. Deuterostomes are in red, mollusks in blue, other Lophotrochozoa in purple, Ecdysozoa in green and Cnidaria in yellow. *Abbreviations: Ac, Aplysia californica; Am, Apis mellifera; Bl, Branchiostoma lanceolatum; Ce, Caenorhabditis elegans; Cg, Crassostrea gigas; Ct, Capitella teleta; Dm, Drosophila melanogaster; Dma, Daphnia magna; Dr, Danio rerio; Es, Euprymna scolopes; Hd, Hybsibius dujardini; Lg, Lottia gigantea; Mm, Mus musculus; Nv, Nematostella vectensis; Ob, Octopus bimaculoides; Ov, Octopus vulgaris; Pd, Platynereis dumerilii; So, Sepia officinalis; Sp, Strongylocentrotus purpuratus; Spo, Schmidtea polychroa; Tc, Tribolium castaneum*.

**Figure S2.2.**
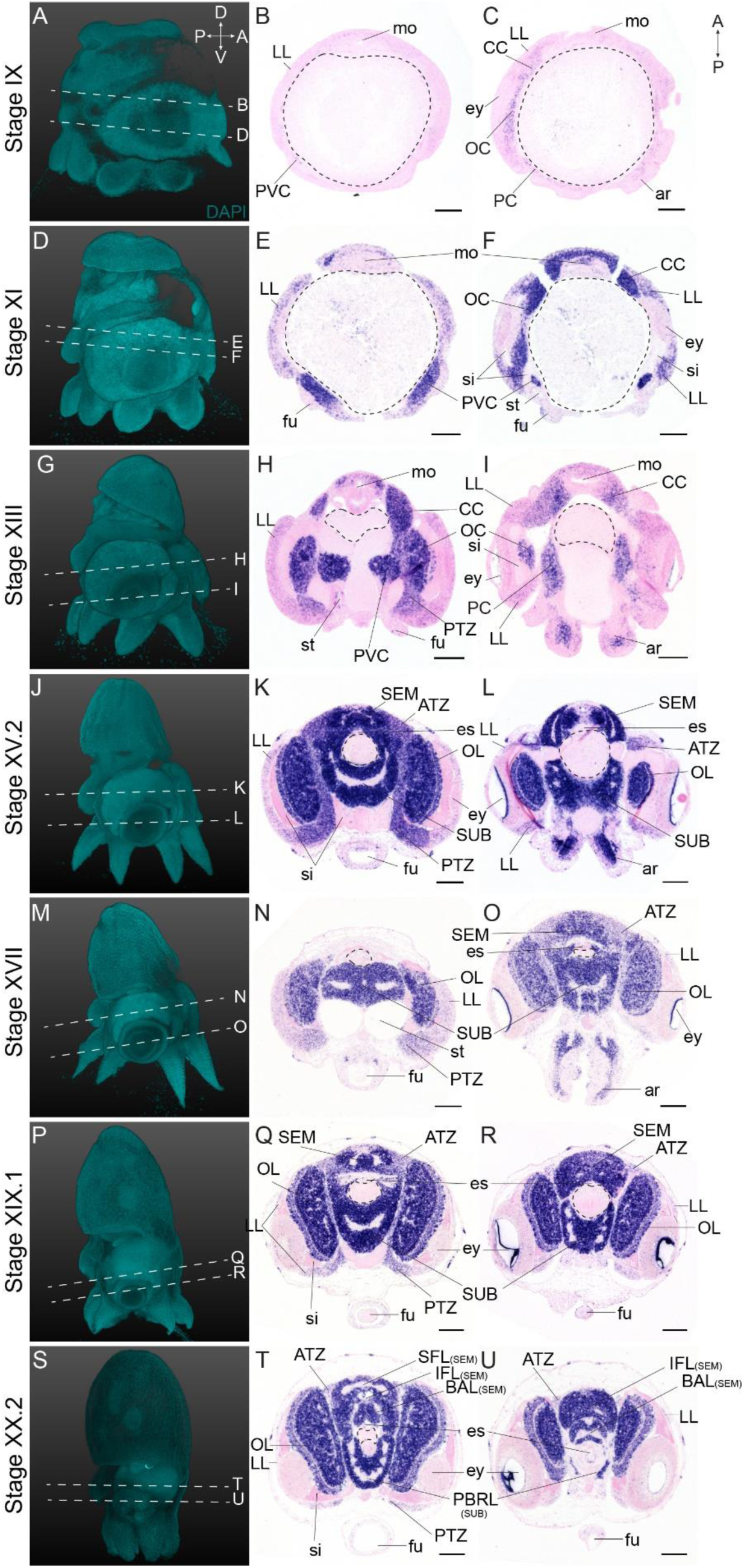
Complementary panels to Figure 2 of *Ov-elav* expression in the head region of developing *O. vulgaris* embryos. Panels on the left show representative 3D reconstructions of embryos seen from the lateral side, stained with DAPI. White dashed lines indicate the sectioning plane in respect to the dorsoventral axis. Panels in the two right columns show *in situ* hybridization of *Ov-elav* on paraffin sections in embryos at Stage IX, XI, XIII, XV.2, XVII, XIX.1 and XX.2, with anterior up and posterior down.

**Figure S2.3.**
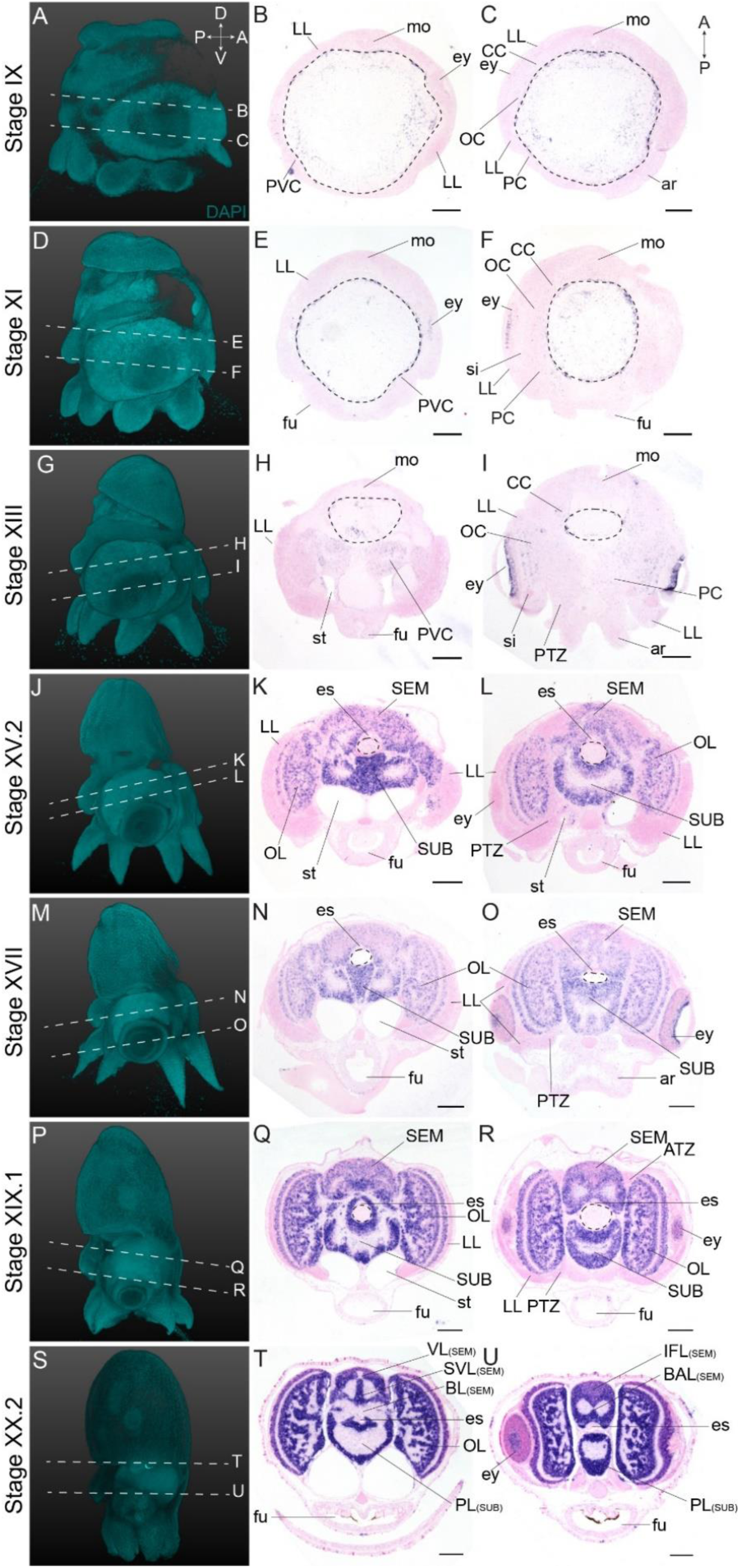
Complementary panels to Figure 2 of *Ov-syt* expression in the head region of developing *O. vulgaris* embryos. Panels on the left show representative 3D reconstructions of embryos seen from the lateral side, stained with DAPI. White dashed lines indicate the sectioning plane in respect to the dorsoventral axis. Panels in the two right columns show *in situ* hybridization of *Ov-syt* on paraffin sections in embryos at Stage IX, XI, XIII, XV.2, XVII, XIX.1 and XX.2, with anterior up and posterior down.

**Figure S4.1.**
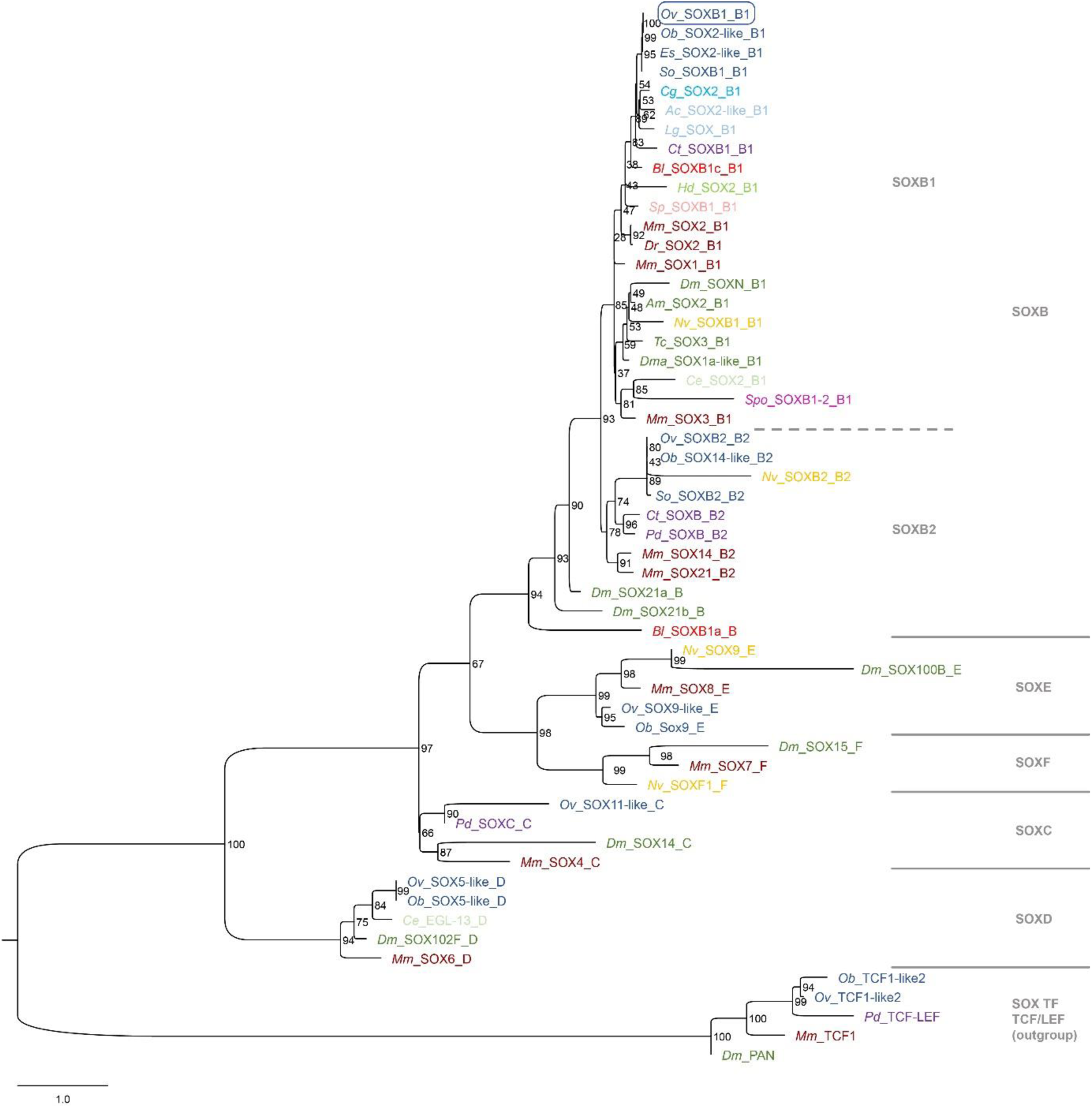
Phylogeny *Ov*-SOXB1. Phylogenetic analysis of SOX family proteins shown in a maximum likelihood tree. Numbers at each branch represent bootstrap support values. The tree has been rooted using the SOX transcription factor family TCF/LEF as outgroup. Sox proteins are divided into 6 families (A-F) with SOXA proteins only present in mammalians and thus excluded from this analysis. Homologs of each group except SOXF could be identified in the available *O. vulgaris* transcriptomes and in other available cephalopod databases. The *O. vulgaris* ID used for subsequent ISH probe design is framed in blue. The scale bar represents the number of amino acid substitutes per site. Deuterostomes are in red, mollusks in blue, other Lophotrochozoa in purple, Ecdysozoa in green and Cnidaria in yellow. *Abbreviations: Ac, Aplysia californica; Am, Apis mellifera; Bl, Branchiostoma lanceolatum; Ce, Caenorhabditis elegans; Cg, Crassostrea gigas; Ct, Capitella teleta; Dm, Drosophila melanogaster; Dma, Daphnia magna; Dr, Danio rerio; Es, Euprymna scolopes; Hd, Hybsibius dujardini; Lg, Lottia gigantea; Mm, Mus musculus; Nv, Nematostella vectensis; Ob, Octopus bimaculoides; Ov, Octopus vulgaris; Pd, Platynereis dumerilii; So, Sepia officinalis; Sp, Strongylocentrotus purpuratus; Spo, Schmidtea polychroa; Tc, Tribolium castaneum*.

**Figure S4.2.**
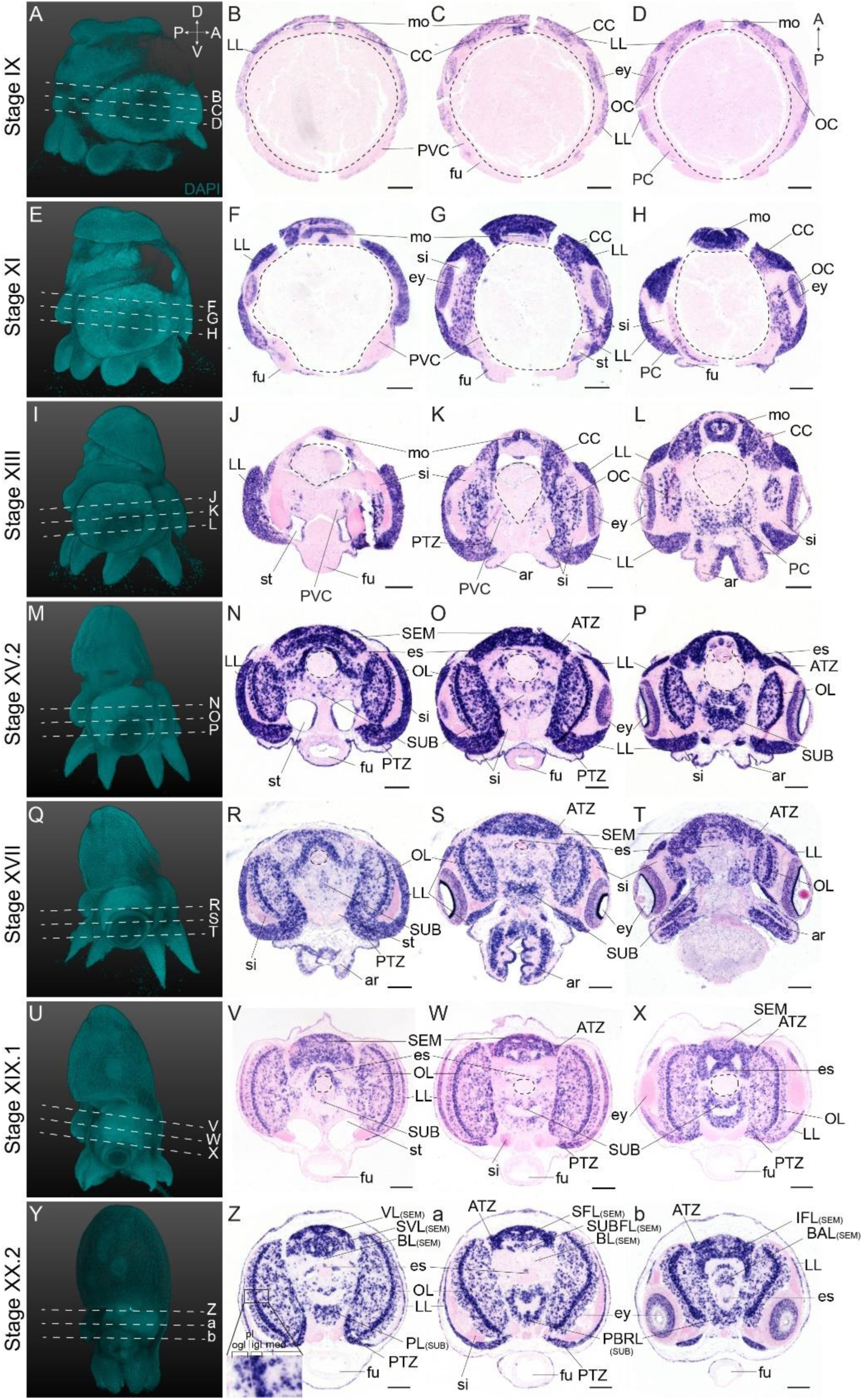
Expression of *Ov-soxB1* during *O. vulgaris* embryonic development. A-b. Panels on the left show representative 3D reconstructions of embryos seen from the lateral side, stained with DAPI. White dashed lines indicate the sectioning plane in respect to the dorsoventral axis. Panels in the three right columns show *in situ* hybridization of *Ov-soxB1* on paraffin sections in embryos at Stage IX, XI, XIII, XV.2, XVII, XIX.1 and XX.2, with anterior up and posterior down. *Ov-soxB1* is expressed in the lateral lips and retina at all stages. In the central brain cords, it is first expressed in the cerebral cord at Stage IX (A-D), is present in the optic and pedal cords from Stage XI onwards (E-H) and appears in the palliovisceral cord at Stage XIII (I-L). *Ov-soxB1* patterns the optic lobes from Stage XV.2 onwards (M-b). *Ov-soxB1* transcripts are also present in the arms, surrounding the nerve cord, and pattern the developing mouth apparatus. Scale bars represent 100 μm. *Abbreviations as in* Figure 1.

**Figure S4.3.**
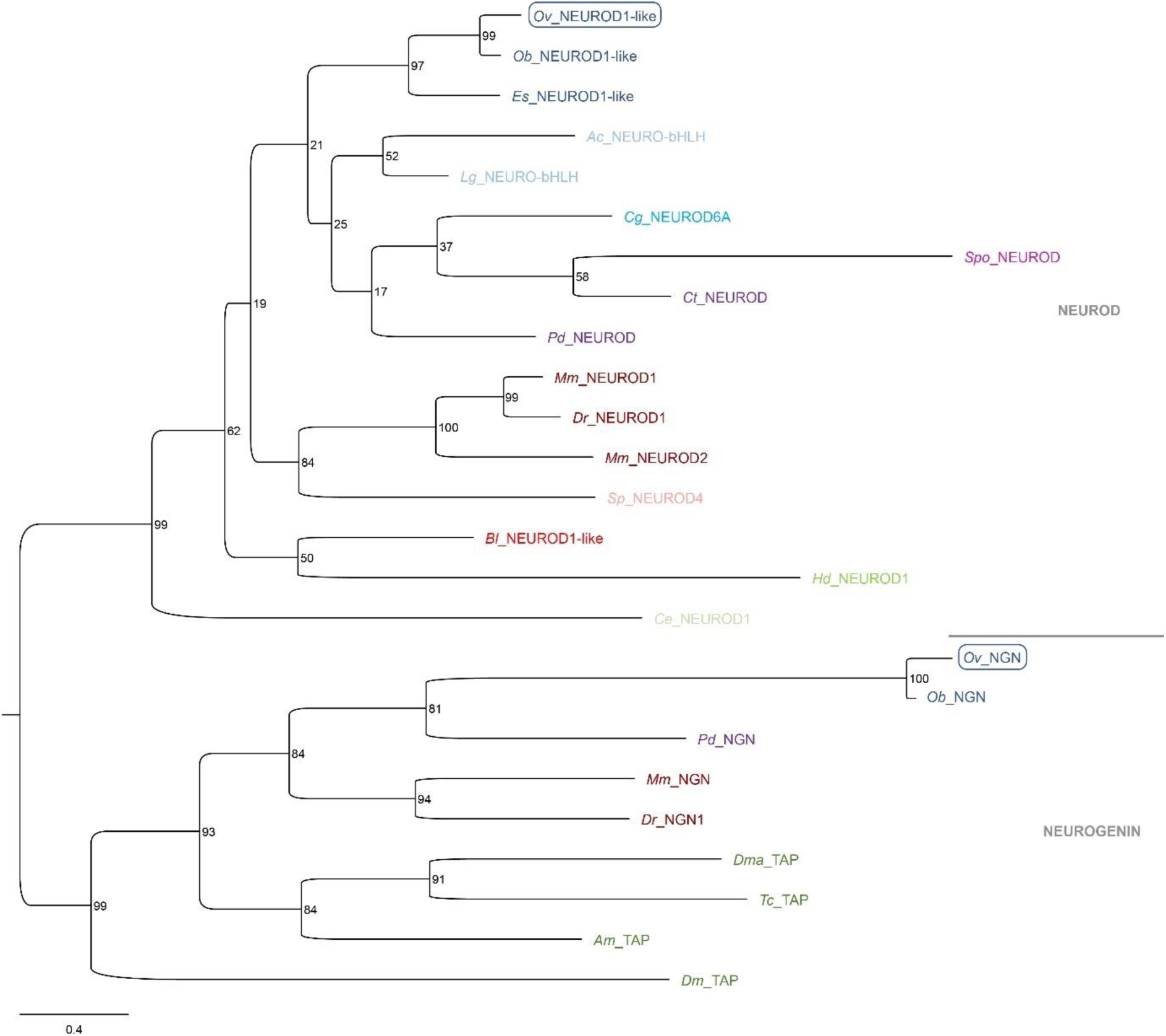
Phylogeny of *Ov-NEUROD* and *Ov-NGN*. A. Phylogenetic analysis of the atonal-related NEUROD, NEUROD-like proteins, NEUROGENIN and TAP proteins shown in a maximum likelihood tree. Numbers at each branch represent bootstrap support values. Arthropods only have a *Neurogenin* homolog *Tap*, but not a *NeuroD* homolog. The *O. vulgaris* IDs used for subsequent ISH probe design are framed in blue. The scale bar represents the number of amino acid substitutes per site. Deuterostomes are in red, mollusks in blue, other Lophotrochozoa in purple, Ecdysozoa in green and Cnidaria in yellow. *Abbreviations: Ac, Aplysia californica; Am, Apis mellifera; Bl, Branchiostoma lanceolatum; Ce, Caenorhabditis elegans; Cg, Crassostrea gigas; Ct, Capitella teleta; Dm, Drosophila melanogaster; Dma, Daphnia magna; Dr, Danio rerio; Es, Euprymna scolopes; Hd, Hybsibius dujardini; Lg, Lottia gigantea; Mm, Mus musculus; Nv, Nematostella vectensis; Ob, Octopus bimaculoides; Ov, Octopus vulgaris; Pd, Platynereis dumerilii; So, Sepia officinalis; Sp, Strongylocentrotus purpuratus; Spo, Schmidtea polychroa; Tc, Tribolium castaneum*.

**Figure S4.4.**
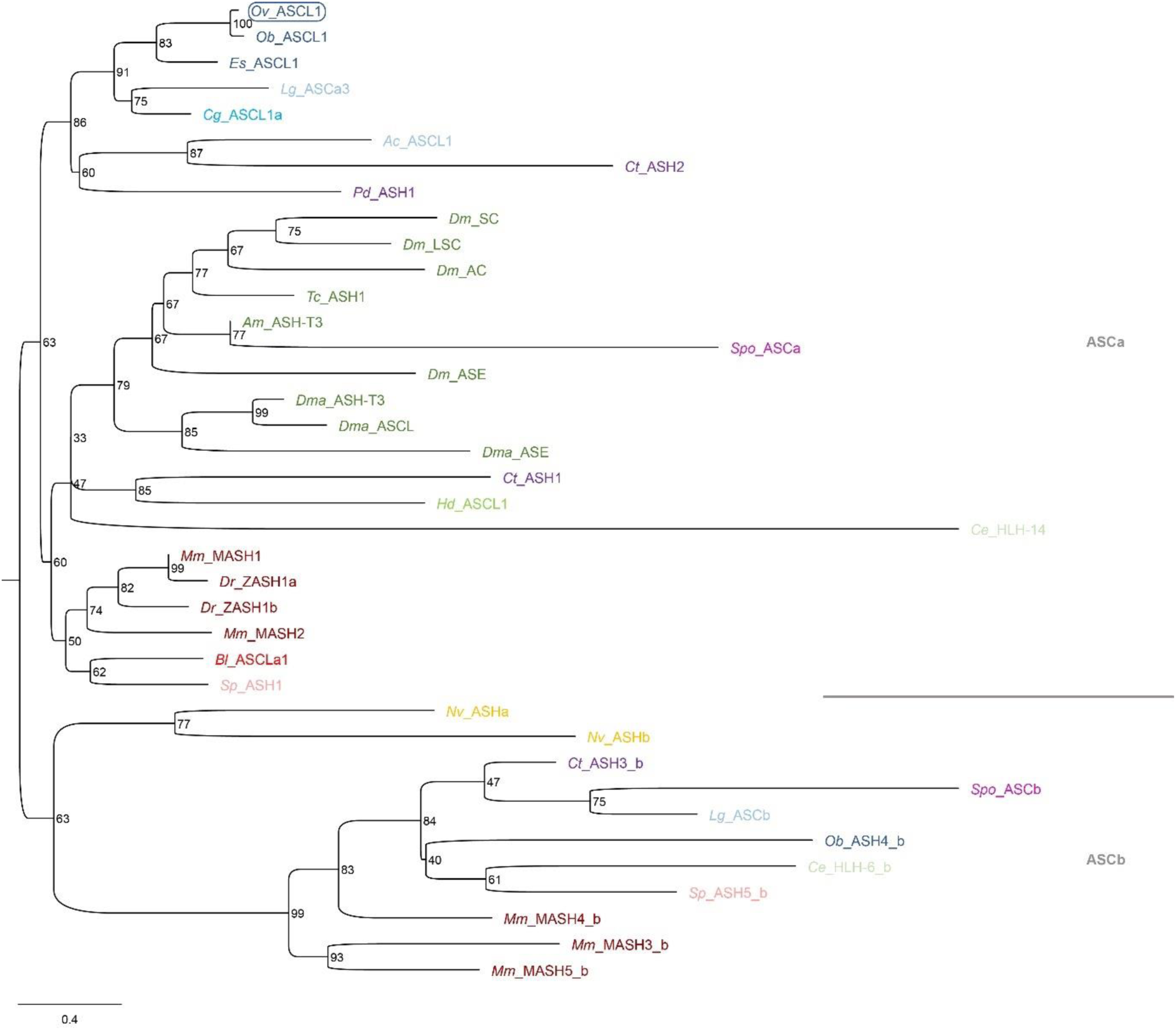
Phylogeny of *Ov-ASCL1*. A. Phylogenetic analysis of ASH and ASH-like proteins shown in a maximum likelihood tree. Numbers at each branch represent bootstrap support values. The tree has been rooted using the achaete-scute ASCb subfamily. The *O. vulgaris* ID used for subsequent ISH probe design is framed in blue. The scale bar represents the number of amino acid substitutes per site. Deuterostomes are in red, mollusks in blue, other Lophotrochozoa in purple, Ecdysozoa in green and Cnidaria in yellow. *Abbreviations: Ac, Aplysia californica; Am, Apis mellifera; Bl, Branchiostoma lanceolatum; Ce, Caenorhabditis elegans; Cg, Crassostrea gigas; Ct, Capitella teleta; Dm, Drosophila melanogaster; Dma, Daphnia magna; Dr, Danio rerio; Es, Euprymna scolopes; Hd, Hybsibius dujardini; Lg, Lottia gigantea; Mm, Mus musculus; Nv, Nematostella vectensis; Ob, Octopus bimaculoides; Ov, Octopus vulgaris; Pd, Platynereis dumerilii; So, Sepia officinalis; Sp, Strongylocentrotus purpuratus; Spo, Schmidtea polychroa; Tc, Tribolium castaneum*.

**Figure S4.5.**
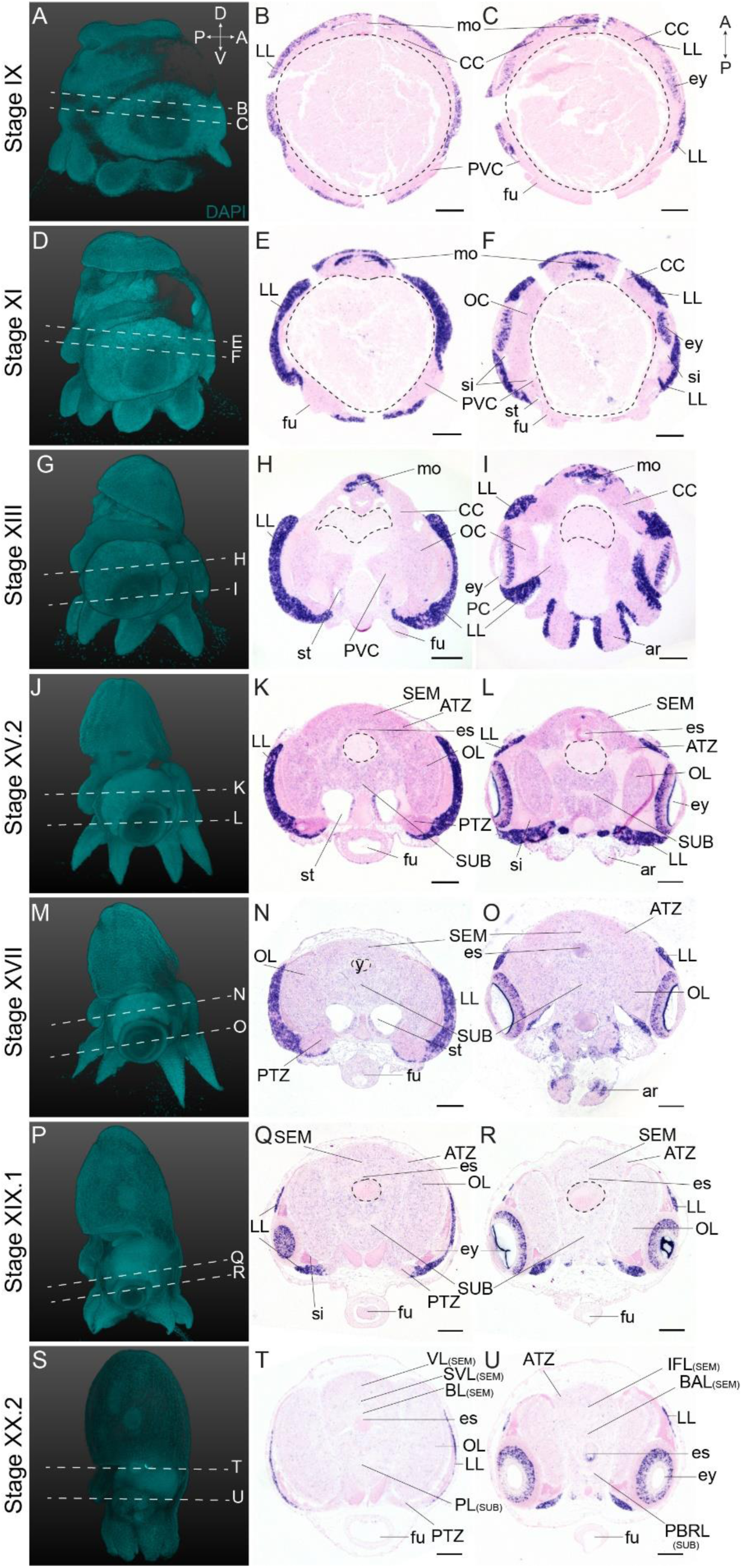
Complementary panels to Figure 4 of *Ov-ascl1* expression in the head region of developing *O. vulgaris* embryos. Panels on the left show representative 3D reconstructions of embryos seen from the lateral side, stained with DAPI. White dashed lines indicate the sectioning plane in respect to the dorsoventral axis. Panels in the two right columns show *in situ* hybridization of *Ov-ascl1* on paraffin sections in embryos at Stage IX, XI, XIII, XV.2, XVII, XIX.1 and XX.2, with anterior up and posterior down.

**Figure S4.6.**
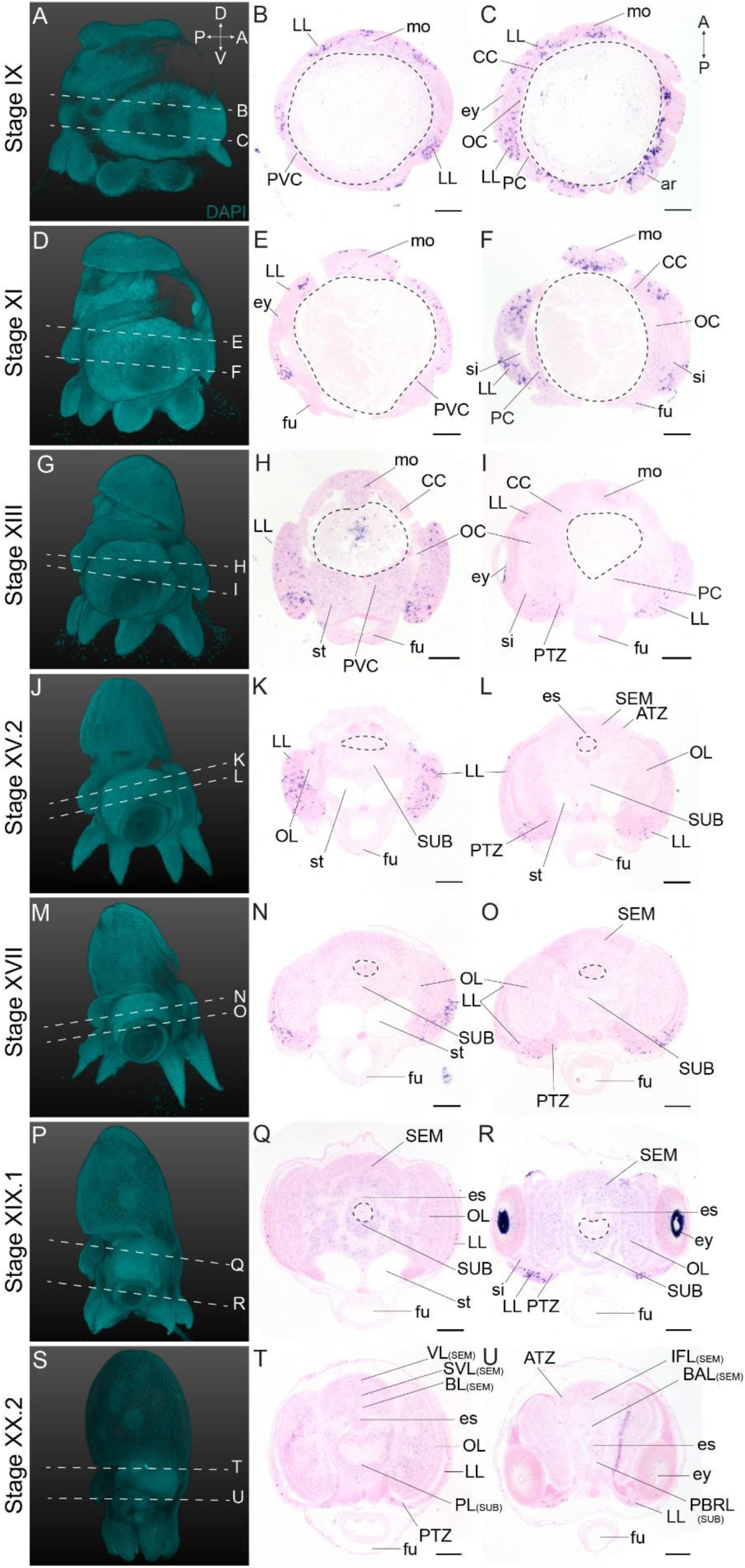
Complementary panels to Figure 4 of *Ov-ngn* expression in the head region of developing *O. vulgaris* embryos. Panels on the left show representative 3D reconstructions of embryos seen from the lateral side, stained with DAPI. White dashed lines indicate the sectioning plane in respect to the dorsoventral axis. Panels in the two right columns show *in situ* hybridization of *Ov-ngn* on paraffin sections in embryos at Stage IX, XI, XIII, XV.2, XVII, XIX.1 and XX.2, with anterior up and posterior down.

**Figure S4.7.**
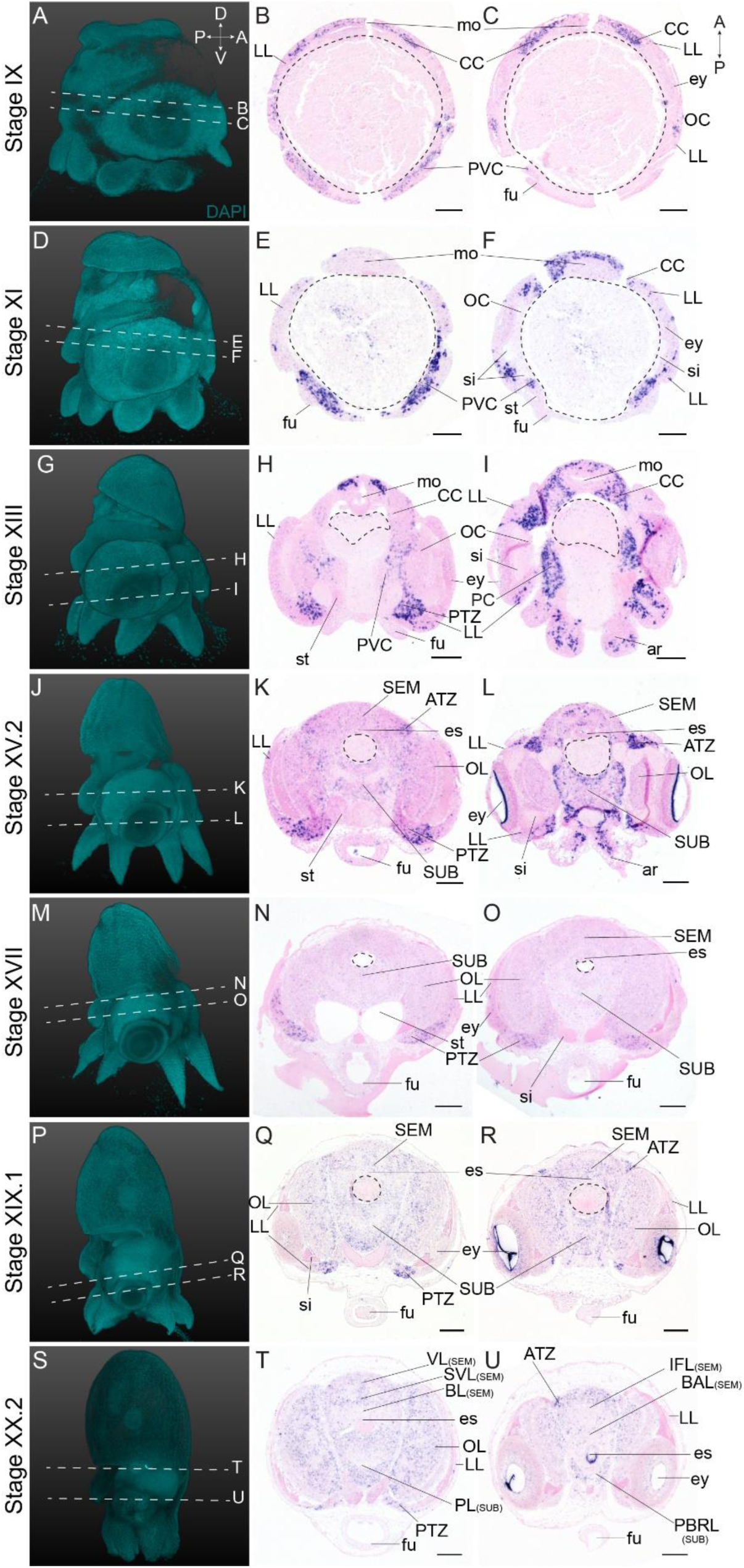
Complementary panels to Figure 4 of *Ov-neuroD* expression in the head region of developing *O. vulgaris* embryos. Panels on the left show representative 3D reconstructions of embryos seen from the lateral side, stained with DAPI. White dashed lines indicate the sectioning plane in respect to the dorsoventral axis. Panels in the two right columns show *in situ* hybridization of *Ov-neuroD* on paraffin sections in embryos at Stage IX, XI, XIII, XV.2, XVII, XIX.1 and XX.2, with anterior up and posterior down.

**Figure S7.1.**
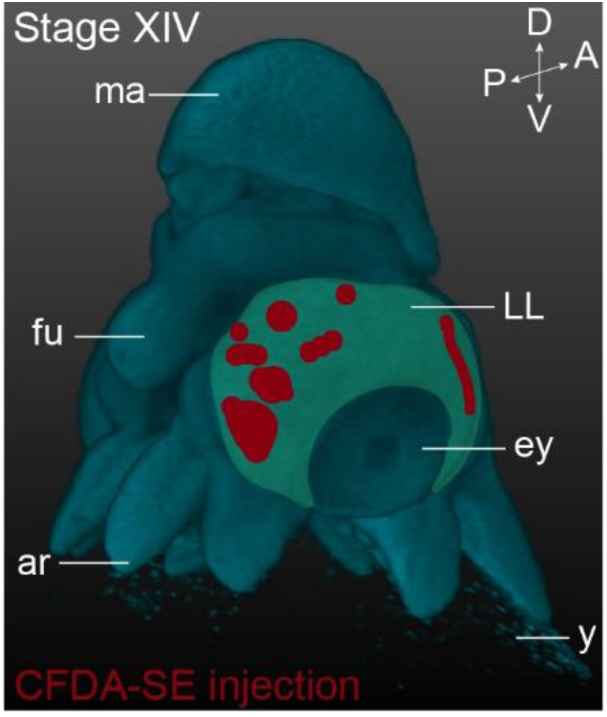
Summary of short-term CFDA-SE lineage tracing. This panel shows the location of the CFDA-SE injection sites at Stage XIV, with each domain representing a single experimental condition. The lateral lips are pseudo-colored in green. *Abbreviations as in* Figure 1.

## Supplementary tables

**Table S1.**
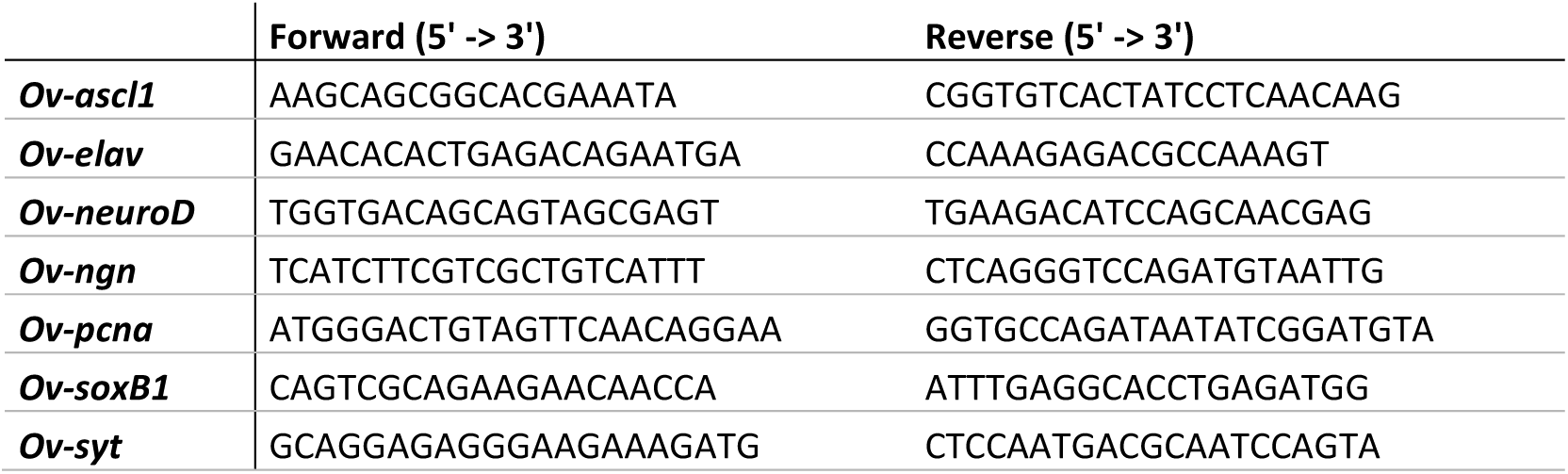
Nucleotide sequence of primers used to amplify gene fragments for ISH probes.

**Table S2.**
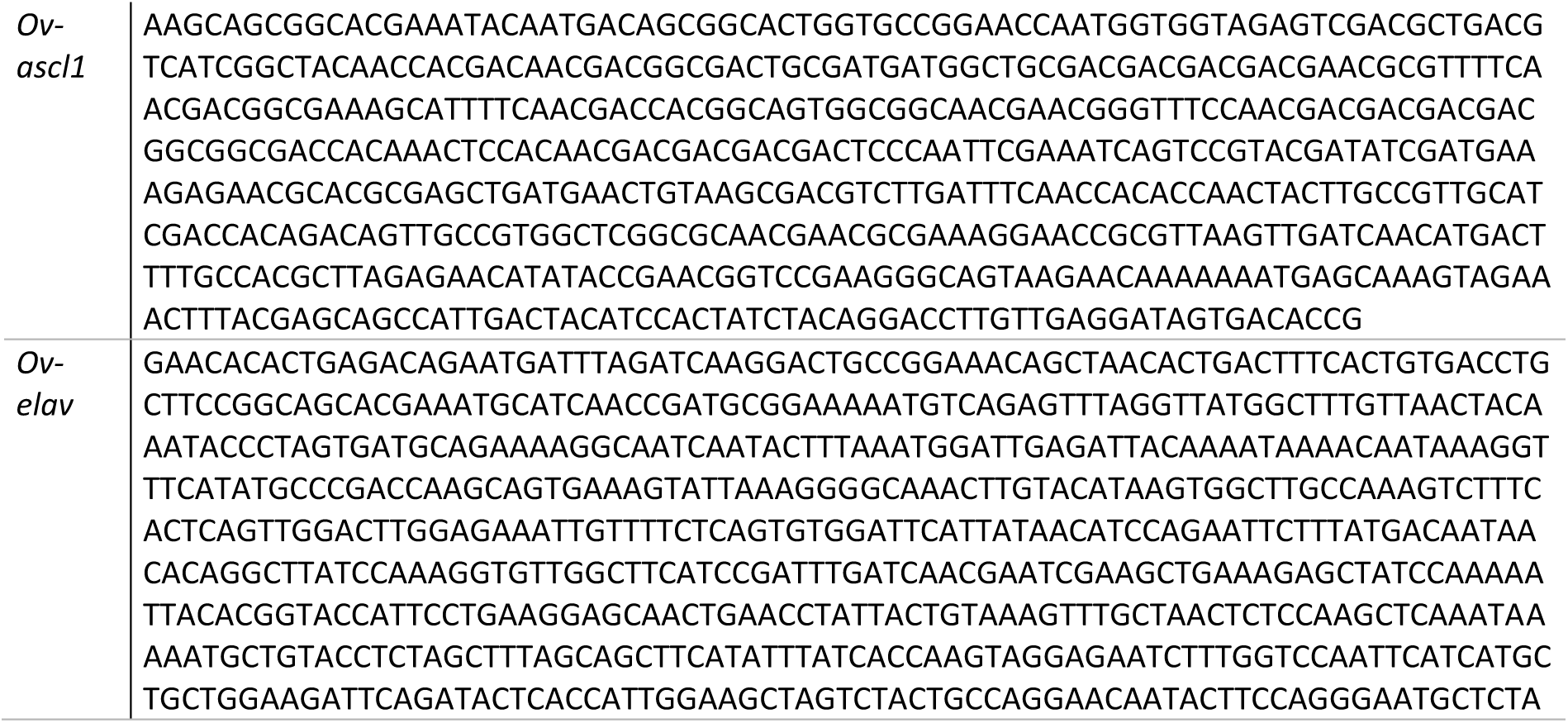
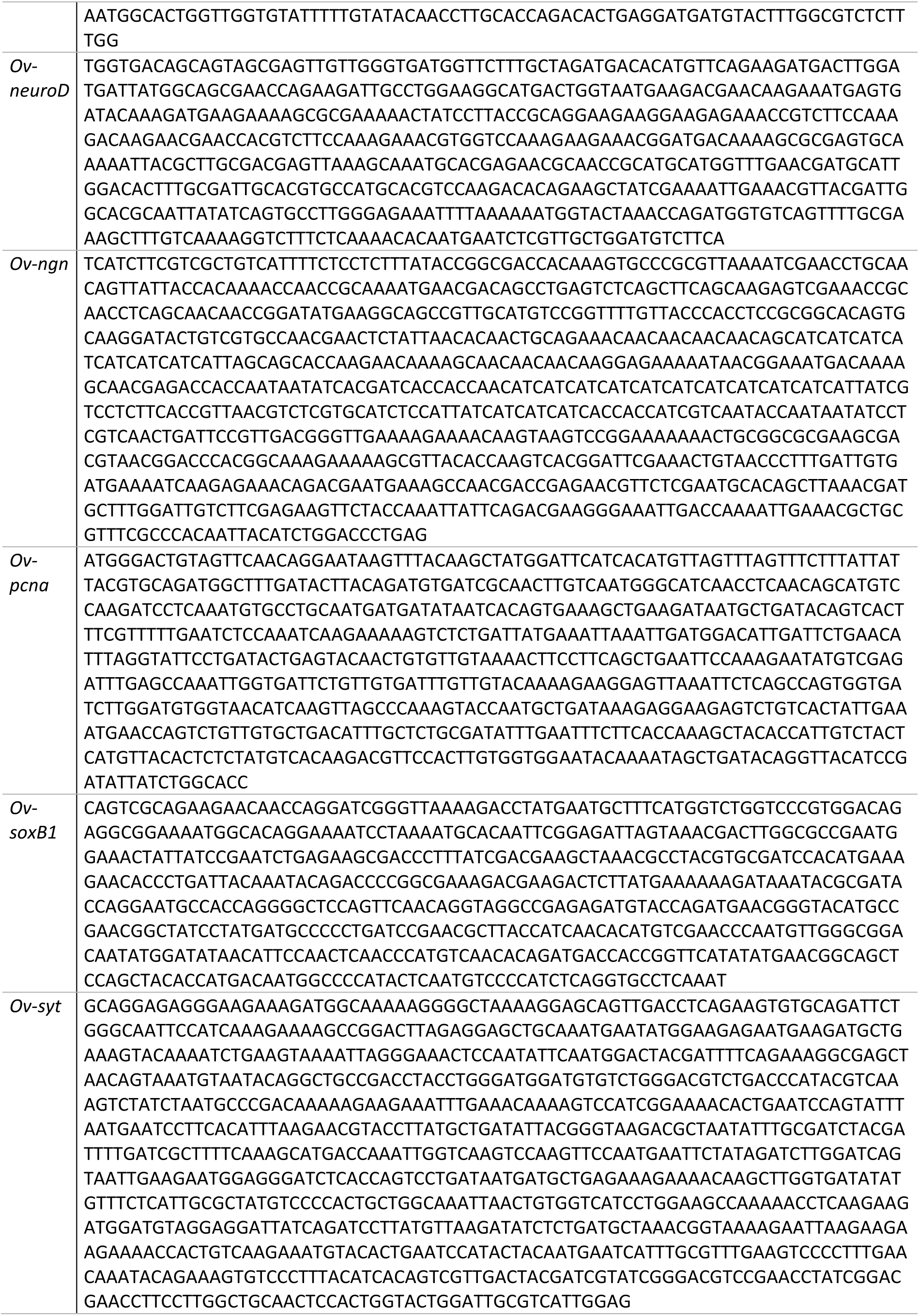
Nucleotide sequence of probes for colorimetric in situ hybridization.

**Table S3.**
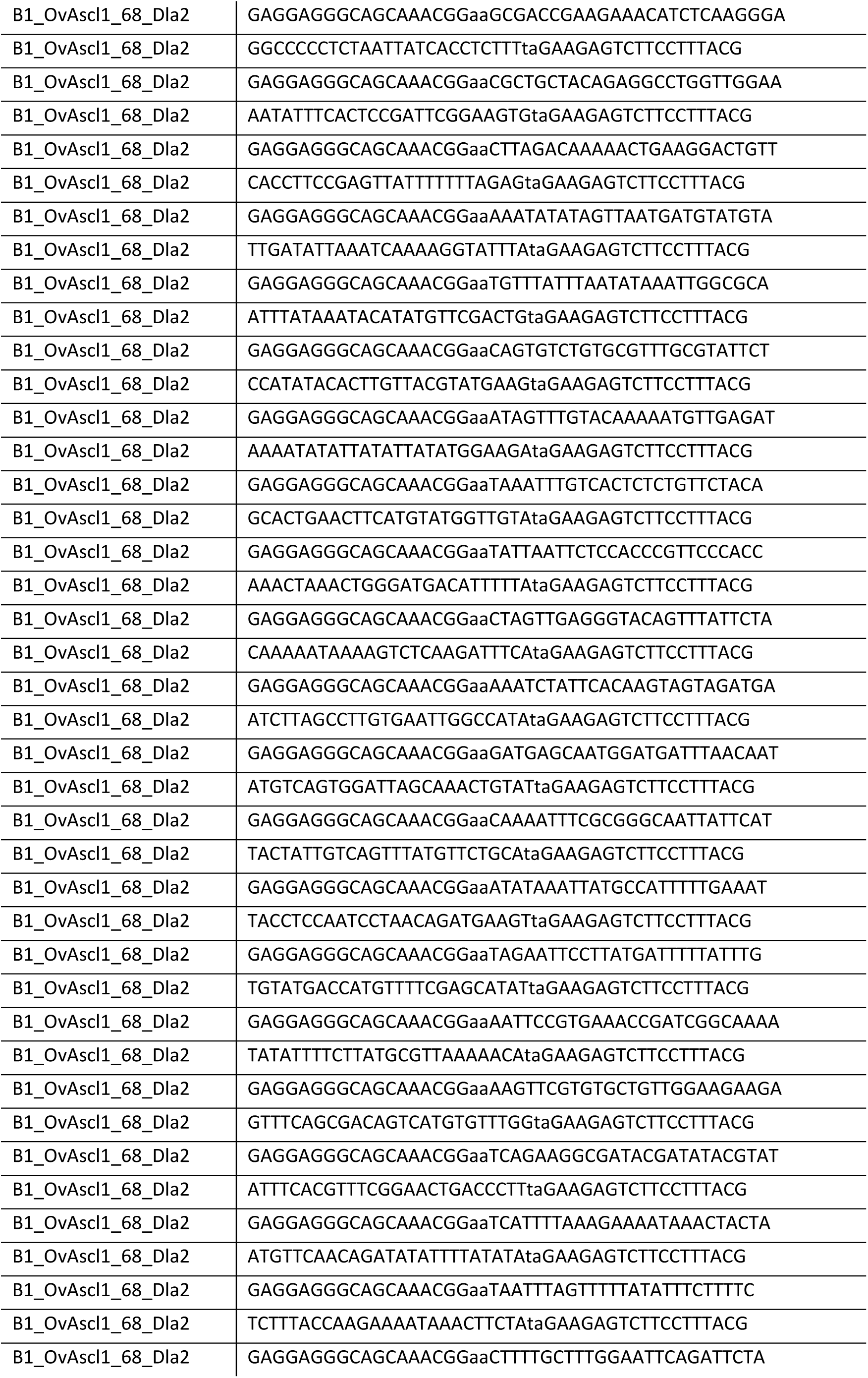
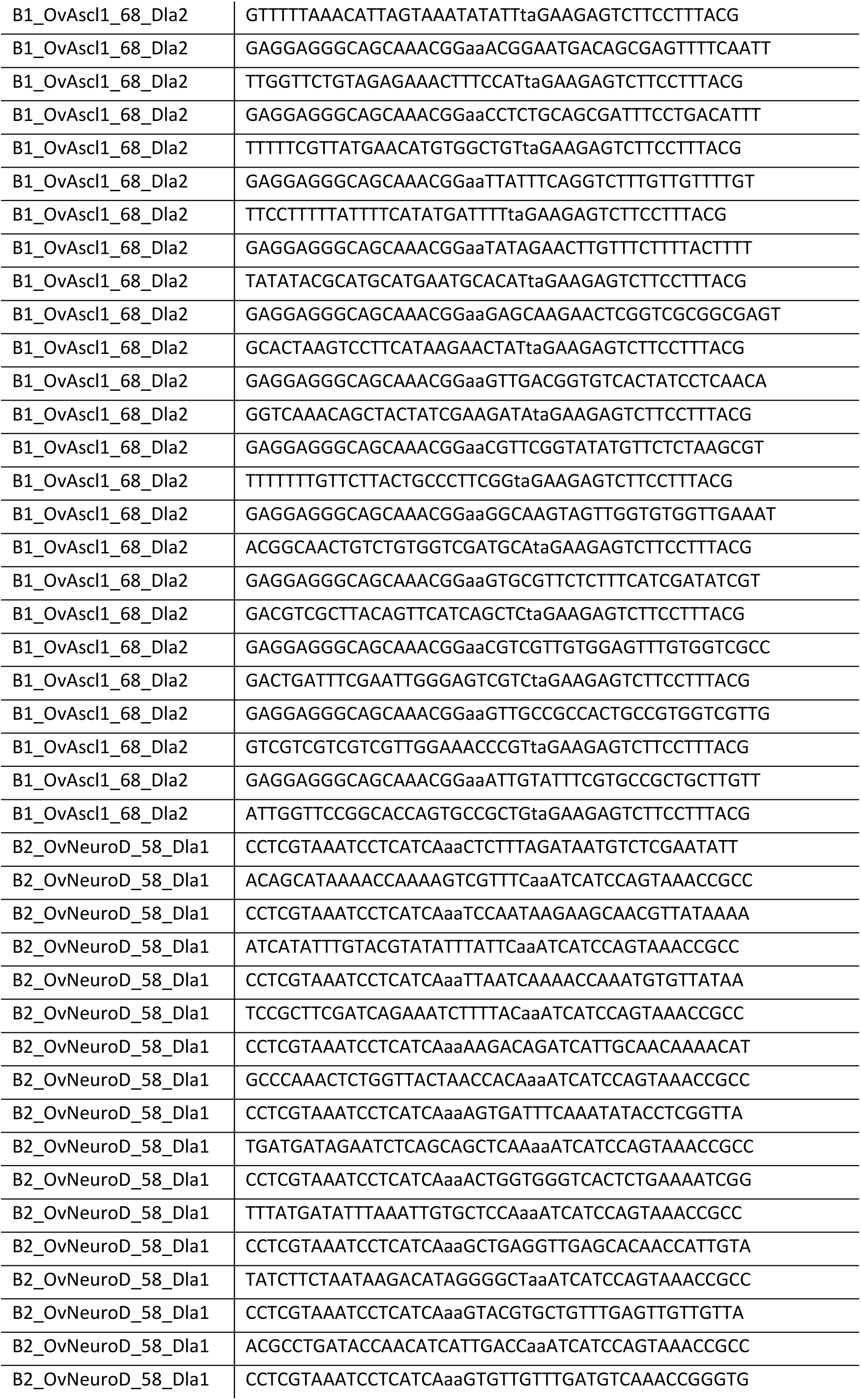
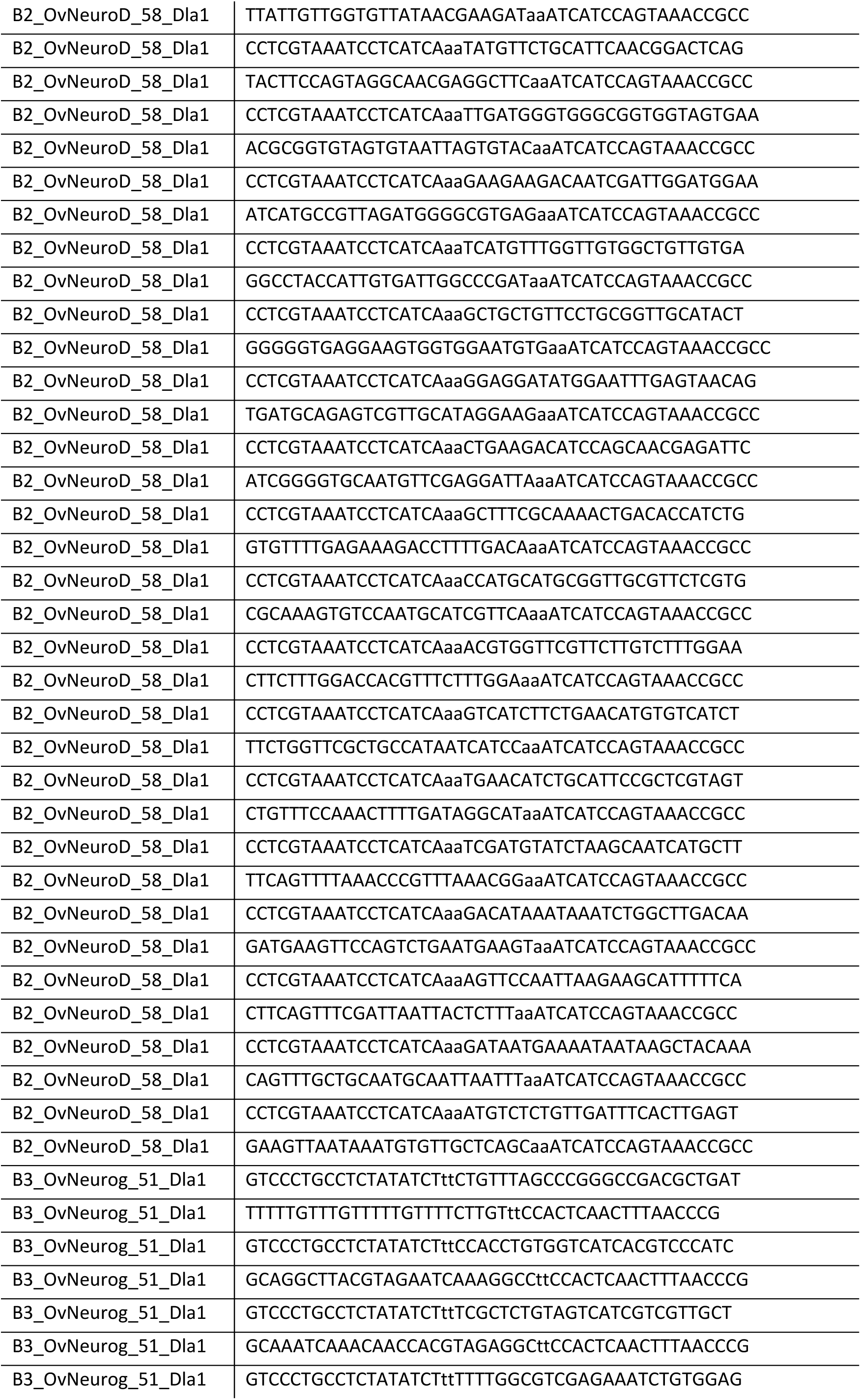
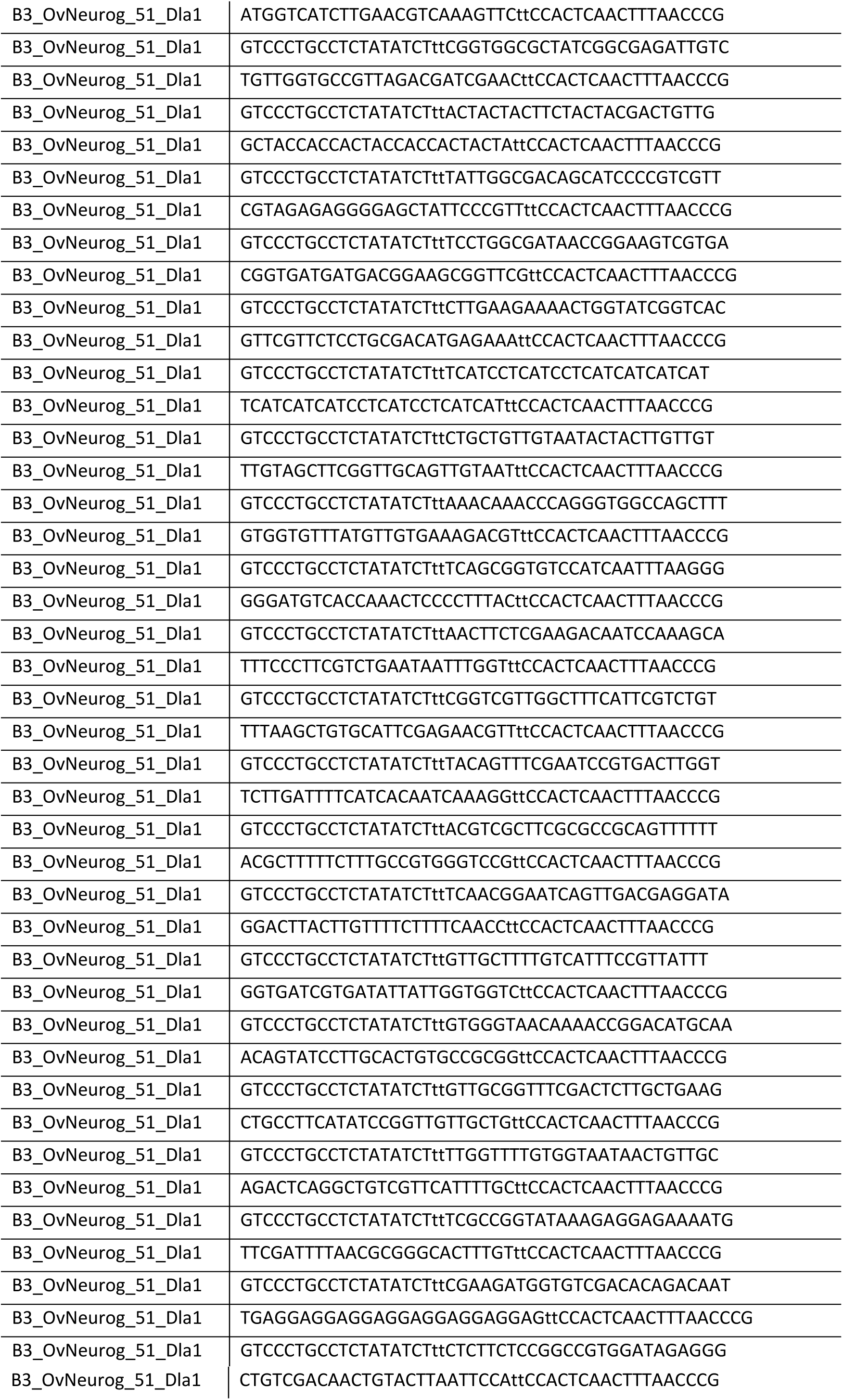
Nucleotide sequence of HCR probes.

**Table S4.** Accession numbers of protein sequences used for phylogenetic tree construction.

| NAME | NCBI accession number | NAME | NCBI accession number |
| --- | --- | --- | --- |
| *Ac-ASCL1* | XP_005110606.1 | *Lg-ASCb* | XP_009064339.1 |
| *Ac-ELAV-like1* | XP_005092587.2 | *Lg-ELAV* | XP_009047849.1 |
| *Ac-ELAV-like2* | AY956294.1 | *Lg-NEURO-bHLH* | XP_009047744.1 |
| *Ac-NEURO-bHLH* | XP_005089777.1 | *Lg-SOX* | XP_009044439.1 |
| *Ac-POLY-A-BINDING4-like* | XP_005113260.1 | *Mm-ELAV-like1* | NP_034615.2 |
| *Ac-SOX2-like* | XP_005093681.1 | *Mm-ELAV-like2* | EDL30947.1 |
| *Am-ASH-T3* | XP_026298936.1 | *Mm-ELAV-like3* | AAH52097.1 |
| *Am-ELAV-like2* | XP_006571248.1 | *Mm-ELAV-like4* | AAH52451.2 |
| *Am-SOX2* | XP_026300481.1 | *Mm-MASH1* | NP_032579.2 |
| *Am-TAP* | XP_001120974.1 | *Mm-MASH2* | NP_032580.2 |
| *Bl-ASCL1a1* | BL13732 (Ensemblmetazoa) | *Mm-MASH3* | NP_064435.1 |
| *Bl-ELAV* | ASW25830.1 | *Mm-MASH4* | NP_001157086.1 |
| *Bl-NEUROD1-like* | BL07636 (Ensemblmetazoa) | *Mm-MASH5* | NP_001257538.1 |
| *Bl-SOXB1a* | ALH22041.1 | *Mm-NEUROD1* | NP_035024.1 |
| *Bl-SOXB1c* | ASW25831.1 | *Mm-NEUROD2* | NP_035025.3 |
| *Ce-EGL-13* | NP_001294841.1 | *Mm-NGN* | AAC52856.1 |
| *Ce-ELAV* | NP_496057.1 | *Mm-POLY-A-BINDING3* | NP_001157308.1 |
| *Ce-HLH14* | NP_495131.3 | *Mm-SOX1* | NP_033259.2 |
| *Ce-HLH6* | CAA87416.1 | *Mm-SOX14* | NP_035570.1 |
| *Ce-NEUROD1* | NP_498115.1 | *Mm-SOX2* | NP_035573.3 |
| *Ce-SOX2* | NP_741836.1 | *Mm-SOX21* | NP_808421.1 |
| *Cg-ASCL1a* | XP_011413561.1 | *Mm-SOX3* | NP_033263.2 |
| *Cg-ELAV-like1* | XP_011437135.1 | *Mm-SOX4* | NP_033264.2 |
| *Cg-NEUROD6A* | XP_011439492.1 | *Mm-SOX6* | AAH67407.1 |
| *Cg-SOX2* | XP_011455662.1 | *Mm-SOX7* | NP_035576.1 |
| *Ct-ASH1* | ACS36111.1 | *Mm-SOX8* | NP_035577.1 |
| *Ct-ASH3* | ELT88014.1 | *Mm-TCF1* | NP_033357.1 |
| *Ct-ELAV1* | ELU12456.1 | *Nv-ASH2* | XP_032226875.1 |
| *Ct-ELAV2* | FJ830867.1 | *Nv-ASH3* | XP_032237760.1 |
| *Ct-NEUROD* | AST23028.1 | *Nv-ELAV* | LT795588.1 |
| *Ct-SOXB* | AST23030.1 | *Nv-SOX9* | XP_001630037.1 |
| *Dma-ASCL* | AEH21241.1 | *Nv-SOXB1* | SJX71971.1 |
| *Dma-ASE* | AEH21242.1 | *Nv-SOXB2* | ABA02364.1 |
| *Dma-ASH-T3* | XP_032780864.1 | *Nv-SOXF1* | ABA02366.1 |
| *Dm-AC* | NP_476824.1 | *Ob-ASCL1* | XP_014781608.1 |
| *Dma-ELAV2* | KZS07880.1 | *Ob-ASH4* | KOF70326.1 |
| *Dm-ASE* | NP_476694.1 | *Ob-ELAV-like1* | XP_014788038.1 |
| *Dma-SOX1a-like* | XP_032796962.1 | *Ob-ELAV-like4* | XP_014767548.1 |
| *Dma-TAP* | XP_032792341.1 | *Ob-NEUROD1-like* | XP_014775193.1 |
| *Dm-ELAV* | P16914.1 | *Ob-NGN* | HM369392.1 |
| *Dm-LSC* | NP_476623.1 | *Ob-POLY-A-BINDING-like1* | XP_014773726.1 |
| *Dm-PAN* | AAC47464.1 | *Ob-SOX14-like* | XP_014789971.1 |
| *Dm*-POLY-A-BINDING | NP_476667.1 | *Ob*-SOX2-like | XP_014780771.1 |
| *Dm*-SC | NP_476803.1 | *Ob*-SOX5-like | XP_014770685.1 |
| *Dm*-SOX100B | NP_651839.1 | *Ob*-SOX9 | XP_014772207.1 |
| *Dm*-SOX102F | NP_726612.1 | *Ov*-TCF1-like2 | XP_014780407.1 |
| *Dm*-SOX14 | NP_476894.1 | *Pd*-ASH1 | CAQ57532.1 |
| *Dm*-SOX15 | NP_523739.2 | *Pd*-ELAV | ABO93208.1 |
| *Dm*-SOX21a | NP_648694.1 | *Pd*-NEUROD | CAQ57533.1 |
| *Dm*-SOX21b | NP_648695.1 | *Pd*-NGN | FM163172.1 |
| *Dm*-SOXN | CAB64386.1 | *Pd*-SOXB | CAY12631.1 |
| *Dm*-TAT | NP_524124.1 | *Pd*-SOXC | CAY12635.1 |
| *Dr*-ELAV-like1 | XP_017213144.1 | *Pd*-TCF/LEF | ANS60442.1 |
| *Dr*-ELAV-like2 | NP_001002172.2 | *So*-ELAV1 | HE956712.1 |
| *Dr*-ELAV-like3 | NP_571524.1 | *So*-ELAV2 | HE956713.1 |
| *Dr*-ELAV-like4 | AAH65965.2 | *So*-SOXB1 | AGL08098.1 |
| *Dr*-NEUROD1 | NP_571053.1 | *So*-SOXB2 | AGL08097.1 |
| *Dr*-NGN1 | NP_571116.1 | *Sp*-ASH1 | XP_003729788.1 |
| *Dr*-POLY-A-BINDING1a | AAH99992.1 | *Sp*-ASH5 | XP_003726868.1 |
| *Dr*-SOX2 | NP_998283.1 | *Sp*-ELAV-like3 | XP_011675251.1 |
| *Dr*-ZACH1a | NP_571294.1 | *Sp*-NEUROD4 | XP_030846597.1 |
| *Dr*-ZACH1b | NP_571306.1 | *Spo*-ASCa | KJ493812.1 |
| *Es*-ASCL | cluster_17772 (Belcaid 2019) | *Spo*-ASCb | AIZ72740.1 |
| *Es*-ELAV-like2/4 | cluster_2695 (Belcaid 2019) | *Spo*-ELAV1 | AIZ76501.1 |
| *Es*-NEUROD1-like | Cluster_3866 (Belcaid 2019) | *Spo*-NEUROD | KJ493814.1 |
| *Es*-SOX2-like | cluster_17675 (Belcaid 2019) | *Spo*-SOXB1-2 | KJ493809.1 |
| *Hd*-ASCL1 | OQV19415.1 | *Sp*-SOXB1 | NP_999639.1 |
| *Hd*-ELAV-like2 | OQV23475.1 | *Tc*-ASH1 | NP_001034537.1 |
| *Hd*-NEUROD1 | OWA52745.1 | *Tc*-ELAV-like3/4 | XP_015833241.1 |
| *Hd*-SOX2 | OQV14806.1 | *Tc*-SOX3 | XP_008193147.1 |
| *Lg*-ASCa3 | XP_009065285.1 | *Tc*-TAP | XP_970244.1 |

**Table S5.**
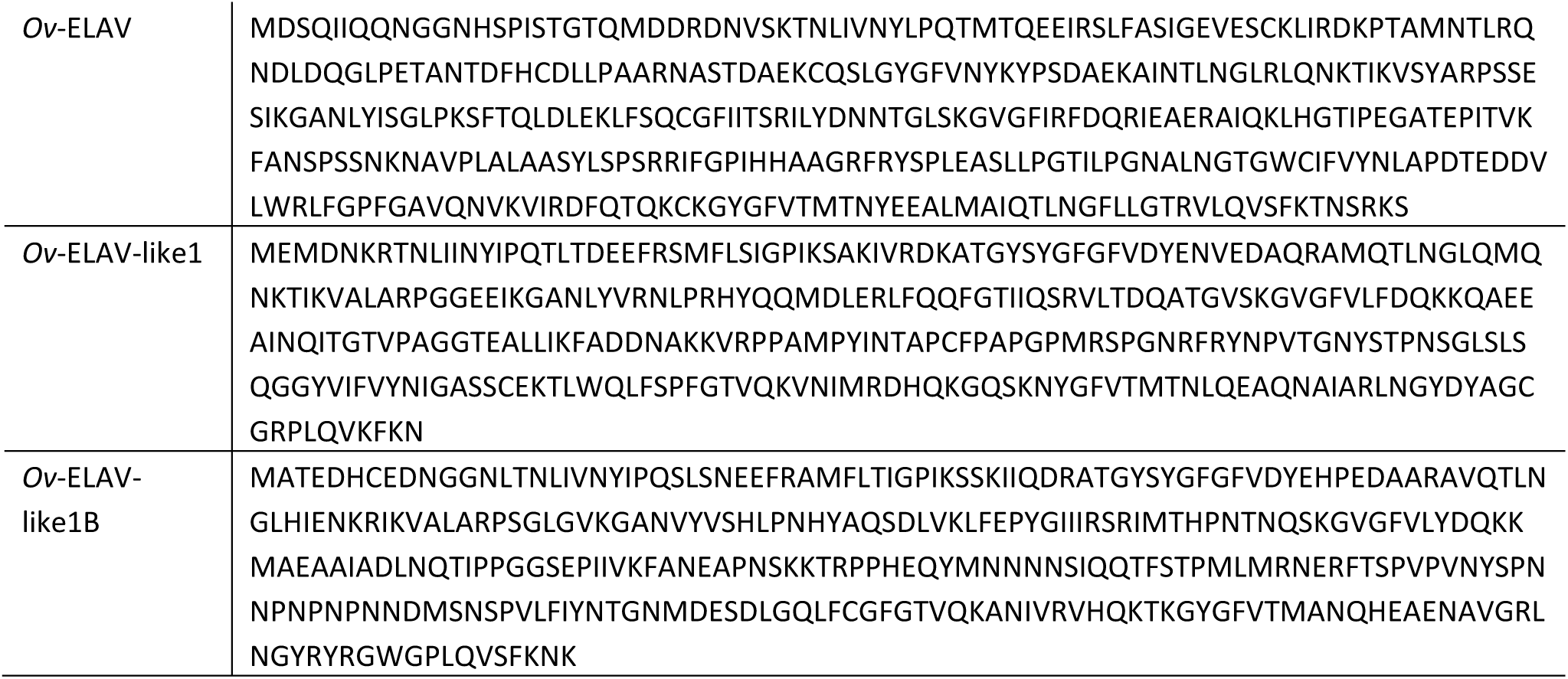
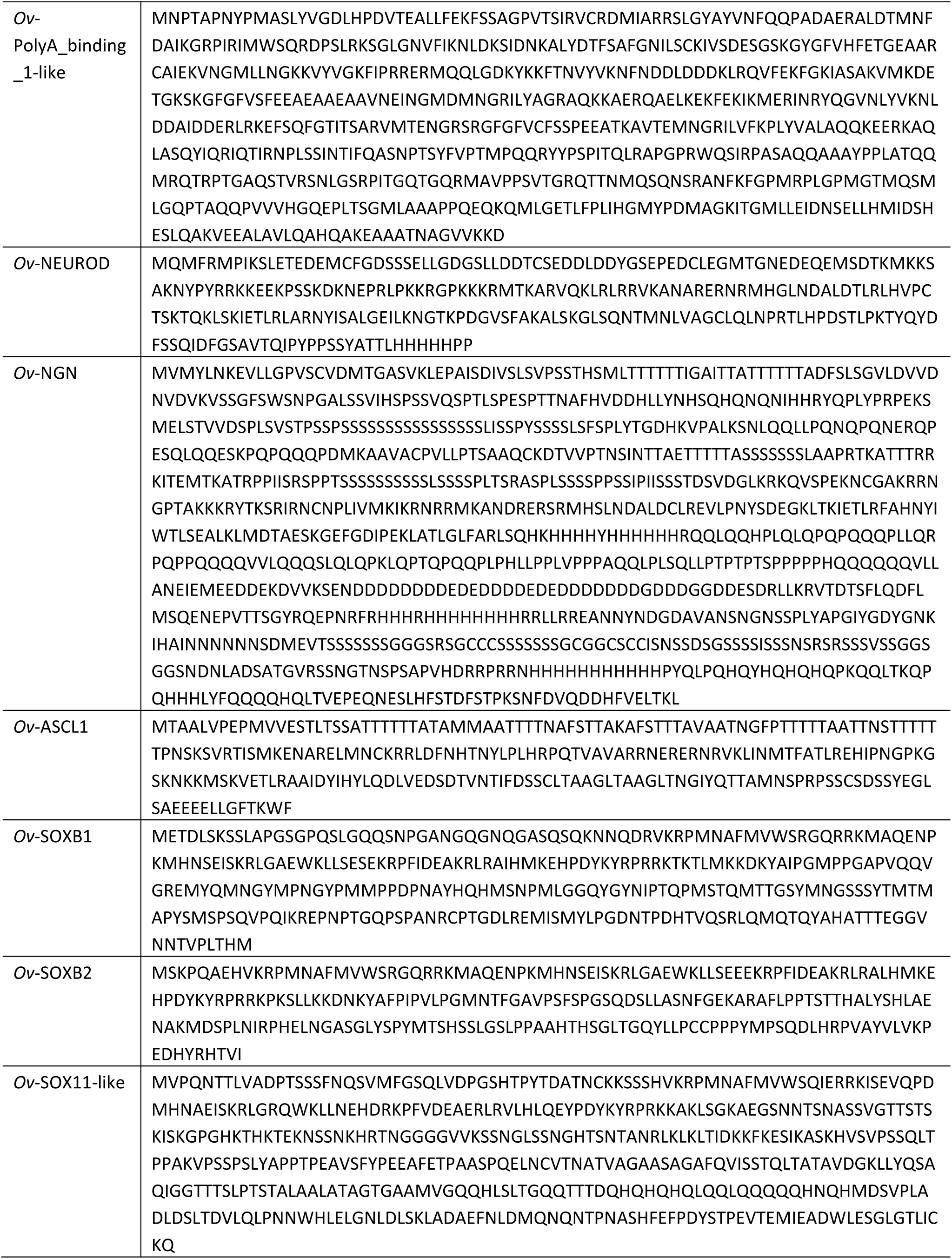
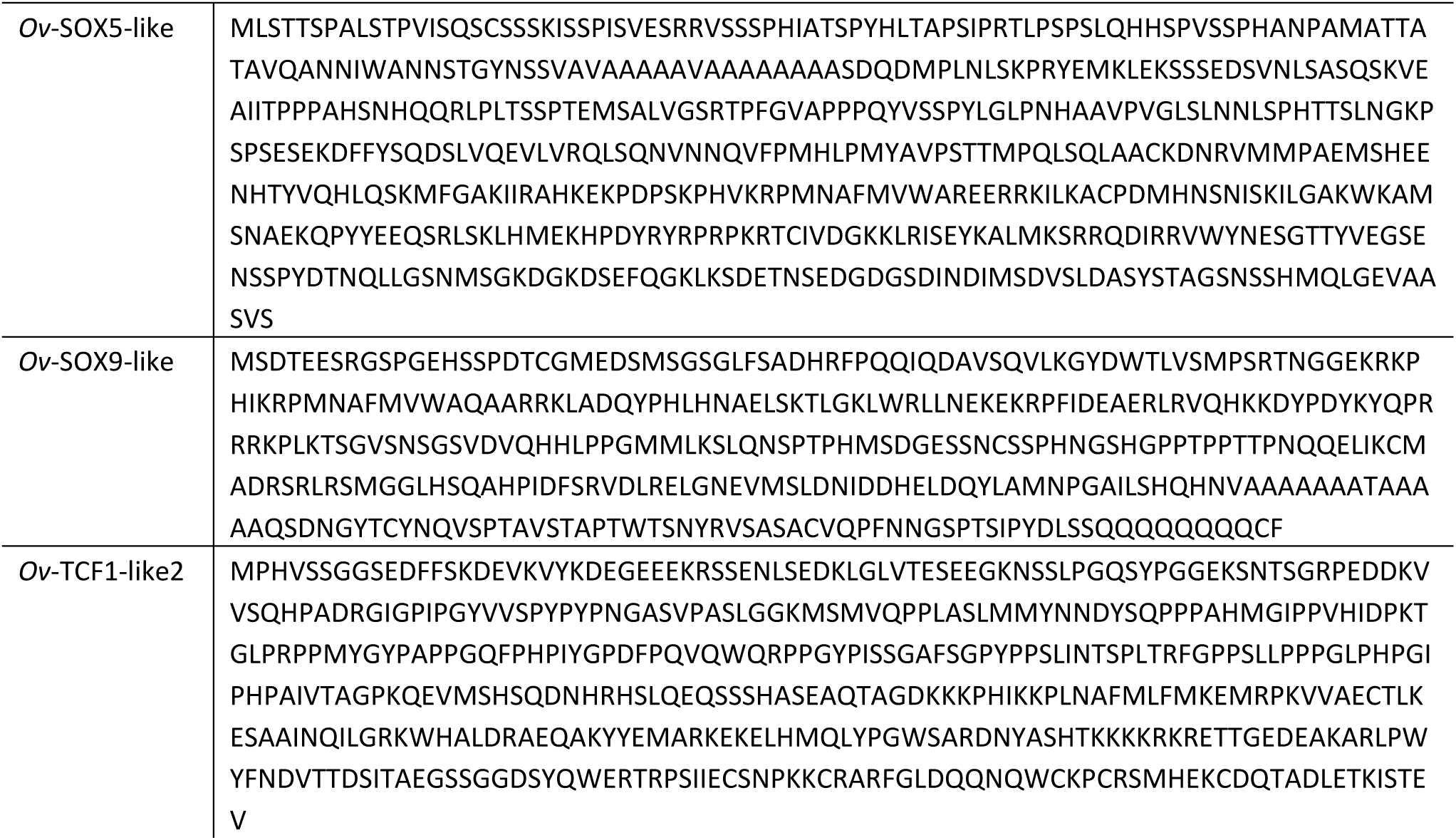
Protein sequences of *O. vulgaris* homologs used in phylogenetic tree reconstruction.

